# Genomic resources of *Ascidiella aspersa* and comparative analysis across tunicates reveal class-level features and evolutionary diversification

**DOI:** 10.1101/2025.11.06.686920

**Authors:** Takumi T. Shito, Vasanthan Jayakumar, Koki Nishitsuji, Yoshie Nishitsuji, Shimon Kawai, Shunsuke O. Miyasaka, Kotaro Oka, Yasubumi Sakakibara, Kohji Hotta

**Affiliations:** Department of Chemistry, Biology and Marine Science, Faculty of Science, University of the Ryukyus, Okinawa, Japan; School of Frontier Engineering, Kitasato University, Sagamihara, Japan; Department of Advanced Aquaculture Science, Faculty of Marine Science and Technology, Fukui Prefectural University, Obama, Fukui, 917-0116, Japan; Department of Bioscience and Informatics, Faculty of Science and Technology, Keio University, Yokohama, 223-8522, Japan; Marine Genomics Unit, Okinawa Institute of Science and Technology Graduate University, Okinawa, Japan

**Keywords:** *De novo* assembly, tunicate, ascidian, Ascidiidae, *Ascidiella aspersa*, codon usage bias, GC content, genome evolution, phylogenomics, evo-devo

## Abstract

**Background:** *Ascidiella aspersa* is an invasive tunicate and one of the closest relatives of vertebrates. Despite its ecological nuisance, *A. aspersa* is anticipated to be a valuable model organism for developmental studies due to its remarkably transparent embryos. However, annotated genome assemblies and transcriptomic resources have not yet been fully established. Although several tunicate genomes have been sequenced, most lack annotations, and a comprehensive analysis across tunicates has not yet been conducted.

**Results:** We performed *de novo* genome assembly and transcriptome analysis of *A. aspersa,* producing a high-quality 306.5 Mb genome assembly. The transcriptome was derived from nine different organs of adults and embryos at six developmental stages. Ab initio and homology-based gene predictions identified 24,504 genes with a BUSCO score of 92.2%. Functional annotation was added for 18,636 genes in the model. To understand the relative features of this species among tunicates, we conducted genome-wide comparative analysis using publicly available data from 35 other tunicate genomes across three classes, five orders, and 12 families, thus constructing gene models for 27 species with BUSCO scores >80%. Overall, phylogenetic analysis revealed a new hypothesis regarding the relationships among Phlebobranchia and Aplousobranchia families. Gene duplication analysis showed distinct contractions of gene families in some taxa with losses of specific DNA repair related genes that were shared among Thaliacea—these may have contributed to their evolutionary diversification. Tunicate genomes exhibited a high level of variation in genomic GC content (28.0%–42.7%). *A. aspersa* has the highest GC content in coding regions and the third position of codons among tunicate species, with changes in codon usage bias differing from other Ascidiidae species. We also constructed an online comparative tunicate genome database (TUNOME), that provides functional annotations of gene models and ortholog analyses based on these genomic and transcriptomic data.

**Conclusions:** We constructed genomic resources for *A. aspersa* and another 35 tunicate gene models. Comparative analysis reveals a variety in tunicate species genomes and characterizes class-level features. Our resources are expected to be a foundation for experimental studies involving non-model tunicates and for comparative analysis among tunicate species.

## Background

Tunicates belong to a sister group of vertebrates and consist of approximately 3,000 species traditionally divided into three classes: Appendicularia, Thaliacea, and Ascidiacea [1]. Their unique evolutionary position makes them important clades in evo-devo studies, particularly in Ascidiacea species, which have sessile adults and planktonic embryos suitable for developmental experiments due to their many unique merits including a short period of embryogenesis, invariant cleavage patterns, and compact genomes [2]. For studying initial development, *Ciona intestinalis* type A (*Ciona robusta*) and *Halocynthia roretzi* are primarily used for studying initial development and have led to significant discoveries in a variety of scientific fields [2, 3].

Recently, *Phallusia mammillata* has emerged as a new model ascidian due to its remarkably transparent embryos, which enable whole embryo live imaging and thereby greatly facilitate the understanding of the developmental mechanisms in chordate [4]. The family Ascidiidae, which includes *P. mammillata*, is characterized by distinctive biological traits such as transparent eggs and embryos [5]. However, as *P. mammillata* is geographically restricted to Europe, alternative model organisms with similar optical transparency are needed in other regions.

One promising candidate is *Ascidiella aspersa,* a cosmopolitan ascidian species belonging to the family Ascidiidae. Although *A. aspersa* is an invasive species that causes biofouling in shellfish aquaculture, particularly in Japan [6–8], its biological characteristics make it an exceptionally valuable experimental organism. The eggs and embryos of *A. aspersa* are highly transparent—transmitting approximately 90% of visible light—and its unfertilized eggs retain mRNA translation capability [5, 9]. Moreover, a standardized developmental table and a three dimensional embryological image resource (R*A*MNE) have already been established for this species [9].

These features, together with its worldwide distribution and ease of collection, highlight *A. aspersa* as a practical and powerful model system for developmental and molecular studies. Although low fertilization efficiency has been a limitation, an improved fertilization protocol has recently been developed to overcome this issue [10]. Despite these advances, a high-quality, annotated genome assembly—an essential foundation for molecular and genetic analyses—has not yet been available for *A. aspersa*.

Since the draft genome of *Ciona intestinalis* [11], many ascidian genomes have been reported. The ANISEED genome database has been developed, which includes model ascidians like *H. roretzi* and *P. mammillata* [3]. ANISEED currently hosts 12 tunicate genomes, and additional tunicate genomes are available in NCBI; however, some lack gene annotation or contain low-quality gene annotations. Furthermore, comprehensive comparative analysis across tunicate genomes has not yet been conducted.

Here, we present a *de novo* genome assembly and transcriptome analysis of *A. aspersa* along with updated gene models for 38 tunicate species. We also compared genomic statistics among species, such as genome size and GC content, thus characterizing specific tunicate taxa. The newly built genomic resources for tunicates will facilitate comparative analysis across tunicate families and expand the use of non-model tunicates in evo-devo studies.

## Methods

### Genome sequencing and assembly of *A. aspersa*

*A. aspersa* specimens were collected from fixed scallop ropes at Onagawa (Onagawa Field Center, Tohoku University) in October 2019. We used sperm as a haploid DNA source to simplify genome assembly and obtain rich amount of DNA, according to the literature [11]. Sperm from a single adult was isolated for DNA extraction. DNA was extracted using the Qiagen DNeasy Kit and quantified using a spectrophotometer and bioanalyzer; subsequently, it was stored at -80°C until sequencing. Hi-C was not performed. The genome sequence was obtained using Pacific Bioscience (PacBio) Sequel and Illumina NovaSeq. PacBio generated over 46.7 Gb of long reads (2.9 million reads), while Illumina paired-end sequencing produced 70 Gb, which was used for three rounds of error correction of long-read data (Table S2).

For genome assembly, wtdbg2 [12], SMARTdenovo [13], Flye [14], and Canu [15] were employed with three rounds of polishing. Repetitive sequences in the genome were identified and masked using Repeat Modeler v2.0.4 [16] and RepeatMasker v4.1.5 [17]. Completeness of the genome assembly was assessed using BUSCO v5 [18] on the gVolante platform [19], utilizing the eukaryota database.

### Transcriptome sequencing and assembly of *A. aspersa*

Samples for the transcriptome analysis were also collected from Onagawa, Hakodate (Hokkaido Research Organization, Hakodate Fisheries Research Institute), and Yoichi (Hokkaido Research Organization, Central Fisheries Research Institute) in Japan. Adult organs were dissected and preserved in RNAlater at -80°C until RNA isolation. Matured eggs isolated from gonoducts were dechorionated using a solution of 0.05% actinase-E and 1% mercaptoacetic acid sodium salt in seawater. Dechorionated eggs and intact embryos were fixed with liquid nitrogen and stored at -80°C. Samples were disrupted by manipulation and repeated freezing with liquid nitrogen, followed by treatment with the RNeasy kit (QIAGEN).

Nine adult organs (gill cage, atrial siphon, oral siphon, fascia, granule, intestinal tract, orange organs, anus, and gonad) were dissected, and eggs or embryos at five developmental stages (gastrula, neurula, early tailbud, mid tailbud, and hatching larva at 25 hpf) were also collected (Fig. S1). For insemination, eggs and sperm were isolated from gonoducts and spermducts, respectively. The sperm were pre-activated with pH 9.5 ASW for 10–15 minutes, following a method used in experiments with *Phallusia* species [20]. Pre-activated sperm were added to eggs in seawater, then diluted fivefold 20 minutes after fertilization. Embryos were incubated at 20°C, and four developmental stages were fixed with liquid nitrogen according to the standard developmental table (Funakoshi et al. 2021). Fixed embryos were stored at -80°C. Embryos with chorions were treated using the TransZol Plant kit (TransGen Biotech).

From the extracted RNA, a stranded library was prepared with polyA selection and paired-end sequencing was performed using Illumina MiSeq and Illumina NovaSeq 6000, producing an average of 62.0 million reads, comprising 8.67 GBases per sample (Table S4). The acquired transcriptome data were quality-checked using FastQC v0.11.9 [21] and SeqKit v2.3.0 [22]. Adapter sequences were trimmed using fastp v0.20.1 [23]. Adapter-trimmed reads were assembled using Trinity v2.14.0 [24]. After adapter trimming, paired-end RNA sequences were aligned to the Trinity assembly using Hisat2 [25] achieving nearly 90% mapping efficiency. Reads that failed to align concordantly were discarded, and the remaining clean reads were subsequently aligned to the genome assembly for downstream analyses (Table S5).

### Gene model construction of tunicate genomes

Gene prediction was performed using two software packages according to Ito et al. [26]. The soft-masked assembly was initially annotated by BRAKER2 pipeline [27] with metazoan proteins of orthoDB10 [28]. To improve prediction accuracy, we also applied the GeMoMa v1.9 pipeline [29], which incorporates RNA-seq evidence and existing gene models of phylogenetically related species for homology-based gene prediction. Gene models from *Ciona robusta* and *Ascidia mentula,* produced with BRAKER2 and having high BUSCO scores, were selected as references for GeMoMa. For exhaustive prediction, genes without start/stop codons were accepted in the GeMoMa step. The two predicted gene models were integrated using GffCompare [30]. We relied on the GeMoMa model, and additional annotations from BRAKER2 were adopted. Translated protein sequences were obtained from the merged gene model using GffRead [30]. We then processed with R scripts to omit alternative isoforms by retaining the longest splicing variant. Functional annotation was added using EnTAP [31] with default settings and the following protein resources: Universal Protein Resource (Swissprot, Trembl) and RefSeq (Invertebrate). BRAKER-predicted genes were filtered by the GeMoMa filter.

Genome sequences were downloaded from repositories such as NCBI or ANISEED. The genome of *Ciona robusta* [32, 33] was downloaded from the Kyoto Ghost Database (https://ghost.zool.kyoto-u.ac.jp/download_ht.html). Genomes of *Ciona savignyi* [34], *Botrylloides schlosseri* [35], *Botrylloides leachii* [36], and *Molgura* spp. [37] as well as *Oikopleura dioica* [38] have been reported and downloaded from ANISEED. *Phallusia mammillata, Phallusia fumigata*, *Halocynthia roretzi*, and *Halocynthia aurantium* have not been published but are available on ANISEED [39]. *Oikopleura dioica* [40], five Oikopleuridae spp. [41], *Salpa aspera*, *Salpa thompsoni* [42], *Clavelina lepadiformis* [43], *Aplidium turbinatum* [44], *Trididemnum clinides* [45], *Corella eumyota* [46], *Corella inflata* [47], *Ascidia mentula* [48], *Styela plicata* [49], and *Styela clava* [50] have been published and are available on NCBI. *Pegea* sp., *Thalia democratica*, *Trididemnum miniatum*, *Trididemnum nubilum*, *Diplosoma virens*, *Perophora annectens*, *Halocynthia papillosa*, and *Boltenia villosa* have not been published but are available on NCBI. First chromosome-level genome assembly of *Botryllus schlosseri* [51] and *Didemnum vexillum* [52] have been published but are unavailable and hence not used in this study. In total, tunicate genomes from 37 species are available in NCBI genome resources, although many of them lack gene models (Table S1).

All available genomes were downloaded. The headers of the FASTA files were processed to be shorter and characters were converted to uppercase before a repeat masking step. For existing gene models, the numbers of genes and transcripts were checked using GeMoMa scripts. BUSCO completeness of protein sequences was assessed using BUSCO v5 on gVolante with the eukaryota dataset. Available gene models with 95% BUSCO value (*Clavelina lepadiformis*, C*iona robusta*, *Styela clava*) were processed with the GeMoMa filter and finalized. Braker2 was performed on other tunicate genomes, while BRAKER3 [53] was used for some genomes (*B. villosa* and *M. erythrocephalus*) due to the failure of BRAKER2 in highly fragmented genomes. Gene models predicted by BRAKER2 with >95% completeness were processed with the GeMoMa filter and finalized. Other genomes with gene models <95% were subsequently processed with GeMoMa gene prediction using high quality gene models from relatively close species. Multiple combinations of one to three reference genomes were tested in each GeMoMa prediction step, and gene models with higher BUSCO values were selected. In three species (*Oikopleura albicans*, *Oikopleura longicauda*, and *Oikopleura vanhoeffeni*), the BRAKER2 alone model was adopted, because it offered more accuracy than integrated models. Alternative splicing was permitted in all prediction steps and processed by R scripts to select the representative longest splicing variants for subsequent analysis. Protein sequences with stop codons were omitted from downloaded models.

### Phylogenetic analysis

OrthoFinder v2.5.4 [54] was used to analyze tunicate amino acid sequences from multiple species’ genomes. One of cephalochordate *Branchiostoma floridae* was selected as an outgroup and genomic data were downloaded from the NCBI database via accession number GCF_000003815.2. Phylogenetic analysis was performed according to Maeda et al [55]. In each orthogroup, gene trees were constructed using MAFFT and FastTree [56] in OrthoFinder. A species tree with all species was drawn from gene trees in orthogroups with all species present using STAG in OrthoFinder [57]. Single-copy orthogroups were used in the following analysis for more precise analysis without the effect of paralogues. Protein sequences were trimmed using trimAl v1.4.rev15 [58] and aligned using MAFFT v7.505 [59]. Aligned sequences were analyzed using IQ-TREE2 v2.1.4-beta [60] for maximum likelihood analysis. Bootstrap resampling with 1,000 pseudo-replicates was performed to obtain statistical support for the nodes. The ML tree was converted to an ultrametric tree based on divergence times between Cephalochordata and the clade comprising Vertebrata and Tunicata (578 Myr) between Stolidobranchia and the clade comprising Thaliacea, Phlebobranchia, and Aplousobranchia (389 Myr); between Thaliacea and the clade comprising Phlebobranchia and Aplousobranchia (296 Myr); and between Molgulidae and the clade comprising Pyulidae and Styelidae (350 Myr) [1]. The conversion was conducted using r8s v1.81 [61] for subsequent analysis.

### Gene duplication analysis

The percentage of duplicated genes was calculated by the ratio of genes assigned to any orthogroup to the total gene number. Rapidly expanded or contracted families were identified using CAFÉ v4.2.1 [62]. For Gene Ontology (GO) analysis, all protein sequences were used as a query to search human protein sequences using BLASTP. Homologies were filtered by BLASTP e-value < 0.01. Genes with the lowest e-value in each orthogroup were used as representative genes. Enrichment analysis used the enrichGO function in the R package clusterProfiler v4.2.1 [63].

### GC content analysis of tunicate genomes

GC content in tunicate genomes was collected from the NCBI database or calculated using SeqKit. Coding sequences (CDS) were extracted using gffread. CDS sequences, including alternative splices, were converted into unique sequences for each locus, and sequences without start/stop codons were omitted using R scripts. The GC content in each position of codons was calculated using R scripts. Statistics were presented using the ggplot2 package in R [64]. Kernel density estimation was performed using the geom_density function. Codon usage of each CDS was calculated using cusp script in EMBOSS [65]. Dimension reduction of codon usage fractions used PCA utilizing the built-in R packages. A parameter evaluating codon usage bias was effective number of codons (ENC) ranging from 20 (only one synonymous codon for encoding amino acids) to 61 (every synonymous codon is used equally) calculated using CodonW v1.4.4 (http://codonw.sourceforge.net/). A smaller ENC value indicates stronger codon usage bias. ENC-GC3 plots and theoretical ENC curves were drawn using the formula (1) [66]. If codon usage bias is affected only by mutation pressure, then the genes will be located on or near the theoretical curve, while natural selection will position them below the curve [67].

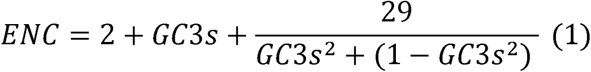

### Genome-wide identification and transcriptomic investigation by TUNOME database

A web-based database of tunicate genomes was constructed using HTML, javascript, and PHP. Functional annotations of gene models for all species were generated by EnTAP. RNA-seq data from different developmental stages of *A. aspersa* were aligned using HiSAT2, and transcript abundance was quantified as TPM using RSEM [68]. The function of genome browser was constructed based on jbrowse2 [69]. These data were integrated to TUNOME database. Gene identification can be performed using BLAST at TUNOME. Gene possession heatmaps were visualized by Rscripts available download page. Colour values were normalized at each gene. Expression heatmaps were visualized by “expression data” page in TUNOME, and values were log2 folded.

## Results

### Genome assembly and transcriptome profiling of *Ascidiella aspersa*

To establish a reference genome for *A. aspersa*, we performed hybrid assembly using long-read (PacBio Sequel) and short-read (Illumina NovaSeq 6000) sequencing achieving approximately 46.8 Gb and 141.4 Gb, respectively. After comparing the results of all four assemblies (Table S3), wtdbg2 assembly—refined through multiple rounds of error correction—was selected for downstream analyses because it yielded the longest contig and the highest N50 value (Table 1). The final assembly had 92.2% BUSCO completeness.

**Table 1.**
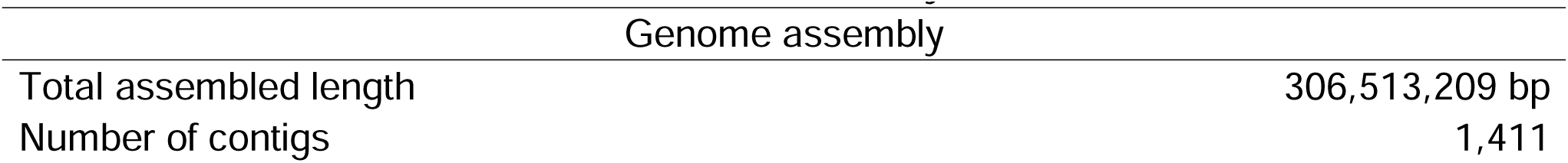
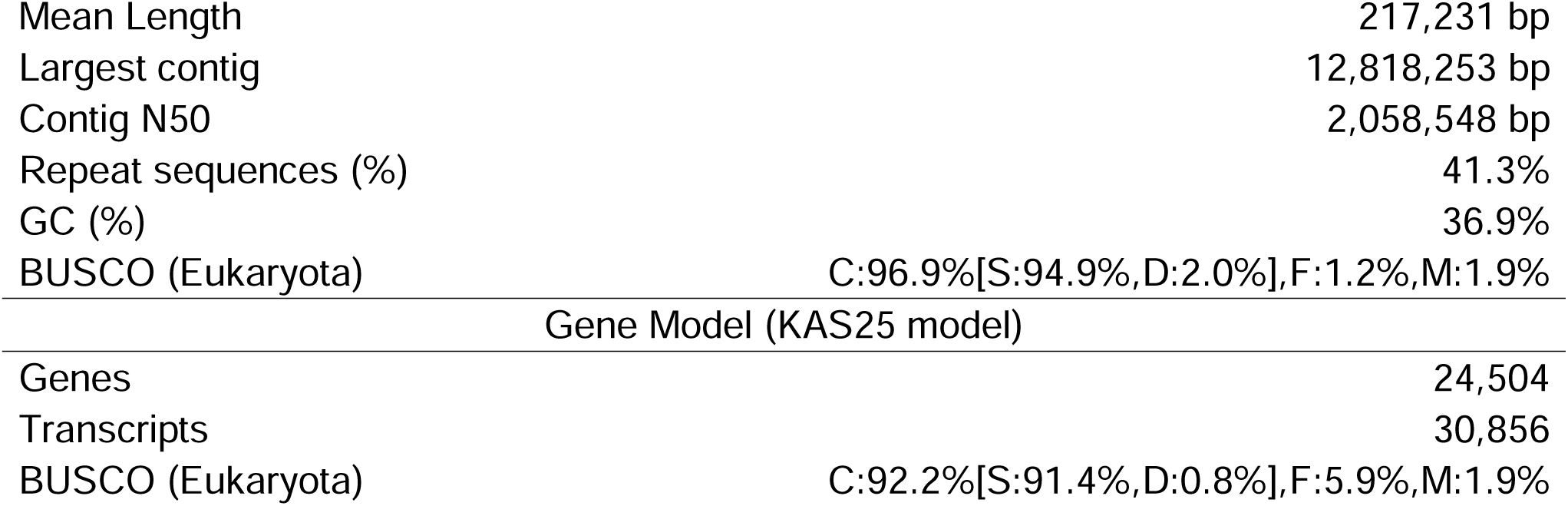
Statistics for the assembly of *A. aspersa*. Statistics of the adopted assembly of the *A. aspersa* genome and constructed gene model (KAS25 model) summary. BUSCO values were assessed on the eukaryote gene sets.

| Genome assembly |  |
| --- | --- |
| Total assembled length | 306,513,209 bp |
| Number of contigs | 1,411 |
| Mean Length | 217,231 bp |
| Largest contig | 12,818,253 bp |
| Contig N50 | 2,058,548 bp |
| Repeat sequences (%) | 41.3% |
| GC (%) | 36.9% |
| BUSCO (Eukaryota) | C:96.9%[S:94.9%,D:2.0%],F:1.2%,M:1.9% |
| Gene Model (KAS25 model) |  |
| Genes | 24,504 |
| Transcripts | 30,856 |
| BUSCO (Eukaryota) | C:92.2%[S:91.4%,D:0.8%],F:5.9%,M:1.9% |

Transcriptome sequencing across nine adult organs and six embryonic stages generated over 60 million reads per sample. The assembled RNA sequences yielded 101,713 transcripts, comprising 97 MBases (Table S5).

Braker2 produced a gene model comprising 20,886 genes and 21,623 transcripts for the *A. aspersa* genome assessed with a relatively low BUSCO score of 69.8% (Table 2). GeMoMa integrated RNA-seq evidence and homology-based refinement predicted 19,302 genes and 25,645 transcripts, thus achieving a higher BUSCO completeness of 92.2% (Table 2; Table S6). Two models were integrated, yielding the final gene model comprises 24,504 genes and 30,856 transcripts with BUSCO completeness of 92.2% (Table 1). This gene model has been designated as the Keio *Ascidiella aspersa* 2025 version (KAS25) gene model.

**Table 2.** Gene prediction and functional annotation of *A. aspersa*. Summary of functional annotations attached to the KAS25 gene model. Gene numbers and transcript numbers containing alternative splicing (BUSCO value) are shown.

|  | Gene/Transcripts<br>(BUSCO) |
| --- | --- |
| Braker2 gene model | 20886/21623 (69.8%) |
| GeMoMa gene model | 19302/25645 (92.2%) |
| Total genes number of KAS25 model | 24504/30856 (92.2%) |
| Total functional annotated gene numbers | 18,636 |
| Total unique sequences with an alignment by similarity search: | 16,872 |
| Total unique transcripts with an alignment: uniprot_sprot | 9,134 |
| Total unique transcripts with an alignment: uniprot_trembl | 14,642 |
| Total unique transcripts with an alignment: refseq_inV | 9,494 |
| Total unique sequences with family assignment: | 18,098 |
| Total unique sequences with at least one GO term: | 15,049 |
| Total unique sequences with at least one pathway (KEGG)<br>assignment: | 5,492 |
| Total unannotated genes | 5,868 |

In total, 18,636 of the 24,504 predicted protein sequences were successfully annotated (Table 2). Homology-based searches identified similarities for 18,636 sequences across the following databases: Uniprot_sprot (9,134 sequences), Uniprot_trembl (14,642 sequences), and Refseq_inV (9,494 sequences). Family-based annotation assigned 18,098 sequences, including 15,049 with GO terms and 5,492 with KEGG terms. In total, 5,868 sequences remained without functional annotation.

### Gene model construction of other tunicate genomes

In total, 37 tunicate genomes spanning three classes, five orders, and 12 families were downloaded from online databases (Table S1). Genome sizes ranged from 64.3 Mb (*Oikopleura dioica*) to 1.1 Gb (*Trididemnum nubilum*). However, BUSCO completeness scores varied widely from 17.3% to 98.9%.

We constructed gene models of 34 species (Table S6). Five species (*Pegea* sp., *Thalia democratica*, *Ascidia mentula*, *Corella eumyota*, and *Botrylloides violaceus*) produced gene models with completeness greater than 95% using BRAKER2, and the remaining 27 species were processed by BRAKER2 and GeMoMa. The number of transcripts—including alternatively spliced isoforms, as well as the BUSCO duplication rate—increased with the number of reference species used. However, the duplication value was generally lower than 10% when alternative isoforms were removed (Table S6).

Overall, our newly constructed gene models showed an average improvement of 15.2% in BUSCO scores compared to 12 existing online models. Finally, 27 of the 34 models we constructed achieved more than 80% BUSCO completeness with the exception of seven species (*Bathochordaeus stygius*, *Mesochordaeus erythrocephalus*, *Oikopleura albicans*, *Oikopleura longicauda*, *Oikopleura vanhoeffeni*, *Boltenia villosa*, and *Botryllus schlosseri*) (Table S7; Table 3).

**Table 3.**
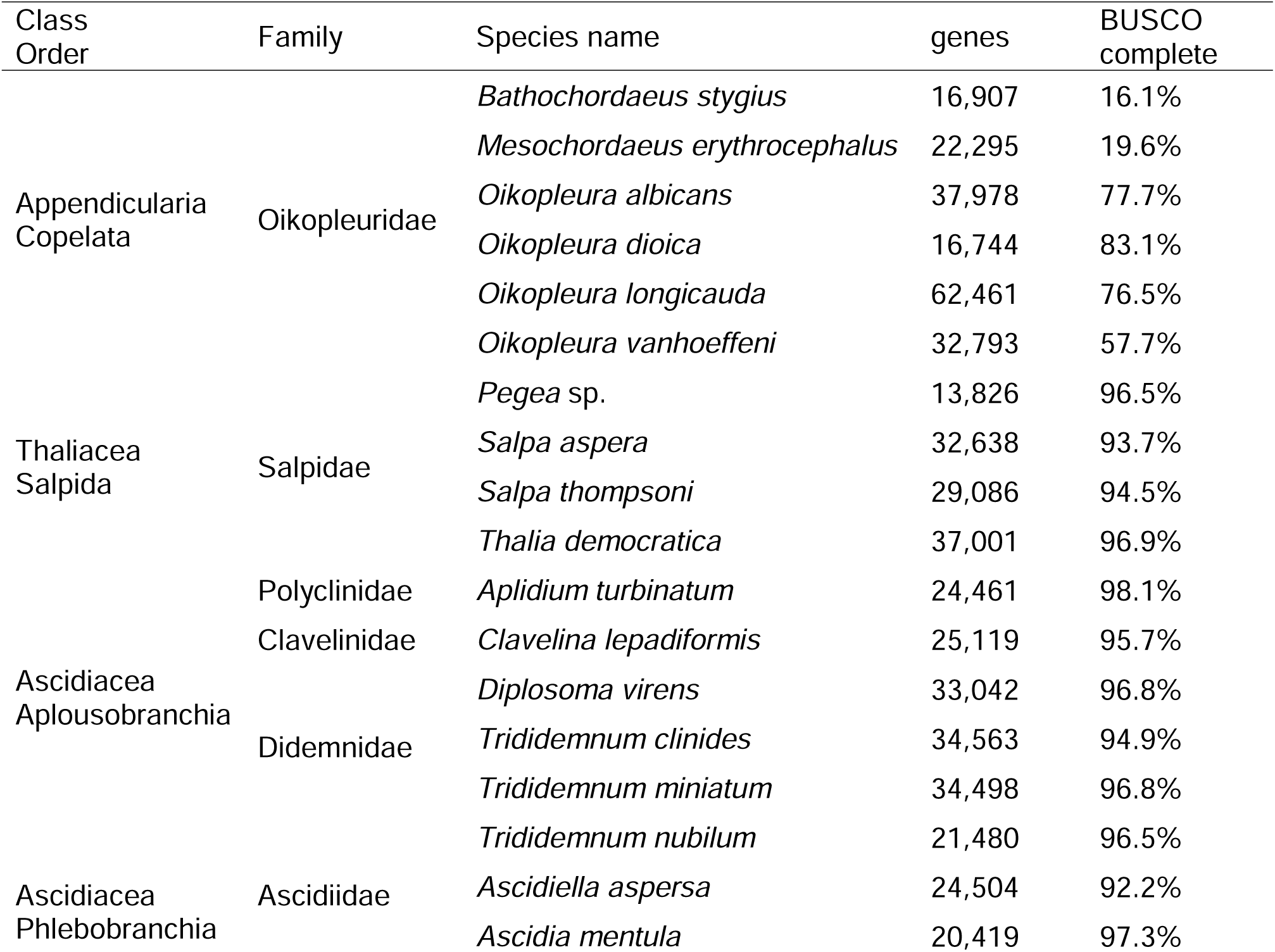
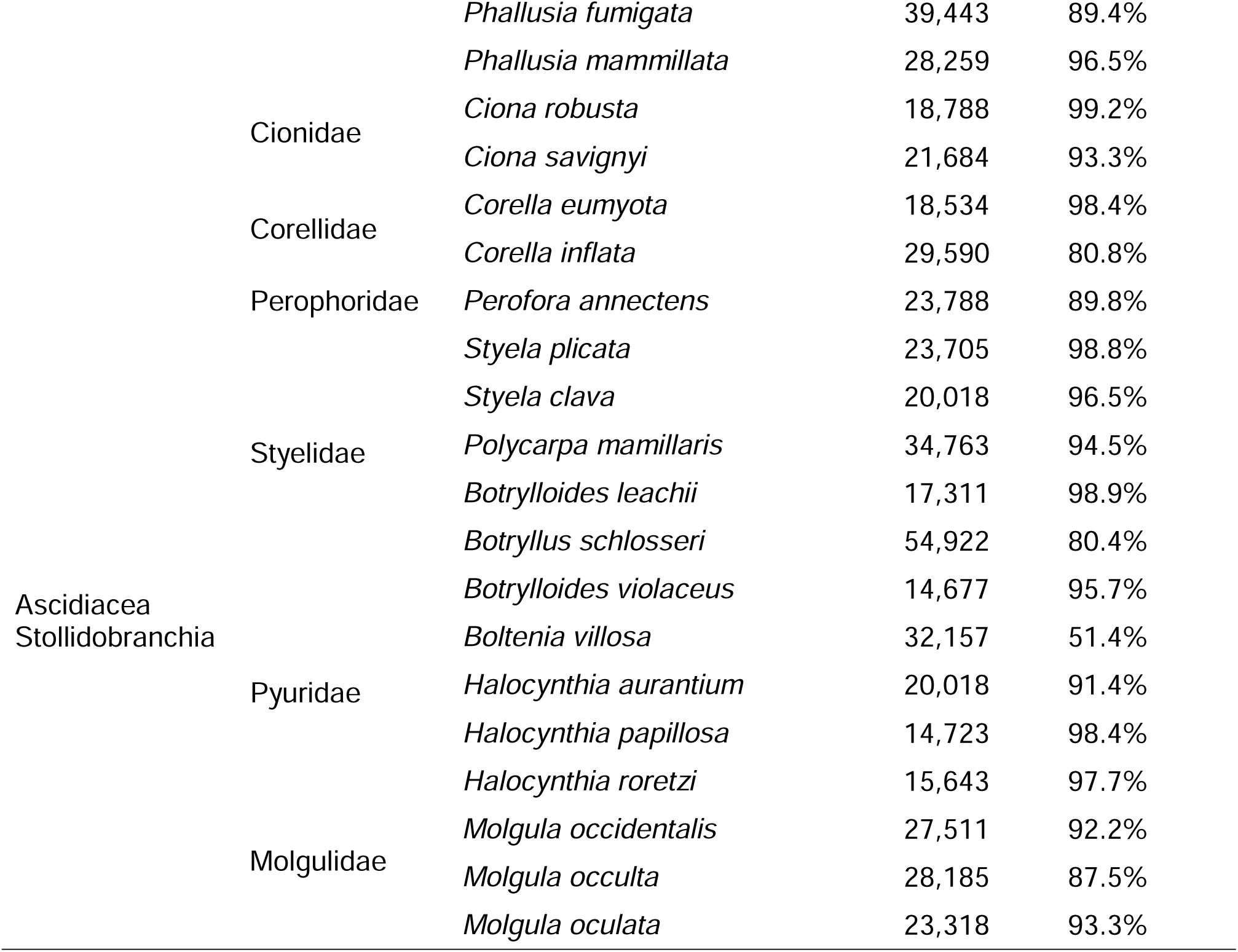
Gene models of all 38 tunicate genomes. Gene numbers, transcript numbers, and BUSCO values of constructed gene models of 38 tunicate genomes. Classes, orders, and families to which they belong are also shown.

### Comparison of gene numbers in constructed gene models

Species with larger genomes generally exhibited a higher number of predicted genes (Fig. S2a–b). *B. schlosseri* and *O. longicauda* exhibit an exceptionally high number of predicted genes relative to their moderate genome sizes compared to other tunicates. Notably, these species show relatively low BUSCO completeness scores (≤80%). Excluding gene models with low BUSCO values, a clearer positive correlation between genome size and gene number was observed.

*Pegea* sp., *B. violaceus*, *H. papillosa*, *H. roretzi*, *O. dioica*, and *B. leachii* have relatively small genomes (<200 Mb) and correspondingly low gene counts (<18,000). There was no strong tendency for closely related species to cluster together in terms of gene number or genome size.

The ratio of duplicated genes to total gene number was also calculated and showed a positive correlation with total gene number (Fig. S2c–d). Species such as *H. papillosa*, *H. roretzi*, *Pegea sp.*, *B. violaceus*, and *O. dioica* exhibited small genomes, low gene counts, and low gene duplication ratios. Since gene duplication rates can be affected by genome completeness, species with BUSCO completeness ≤90% were excluded from further analysis. This exclusion clarified the trend: species with larger genomes tended to have higher numbers of duplicated genes (Fig. 1a).

**Fig. 1.**
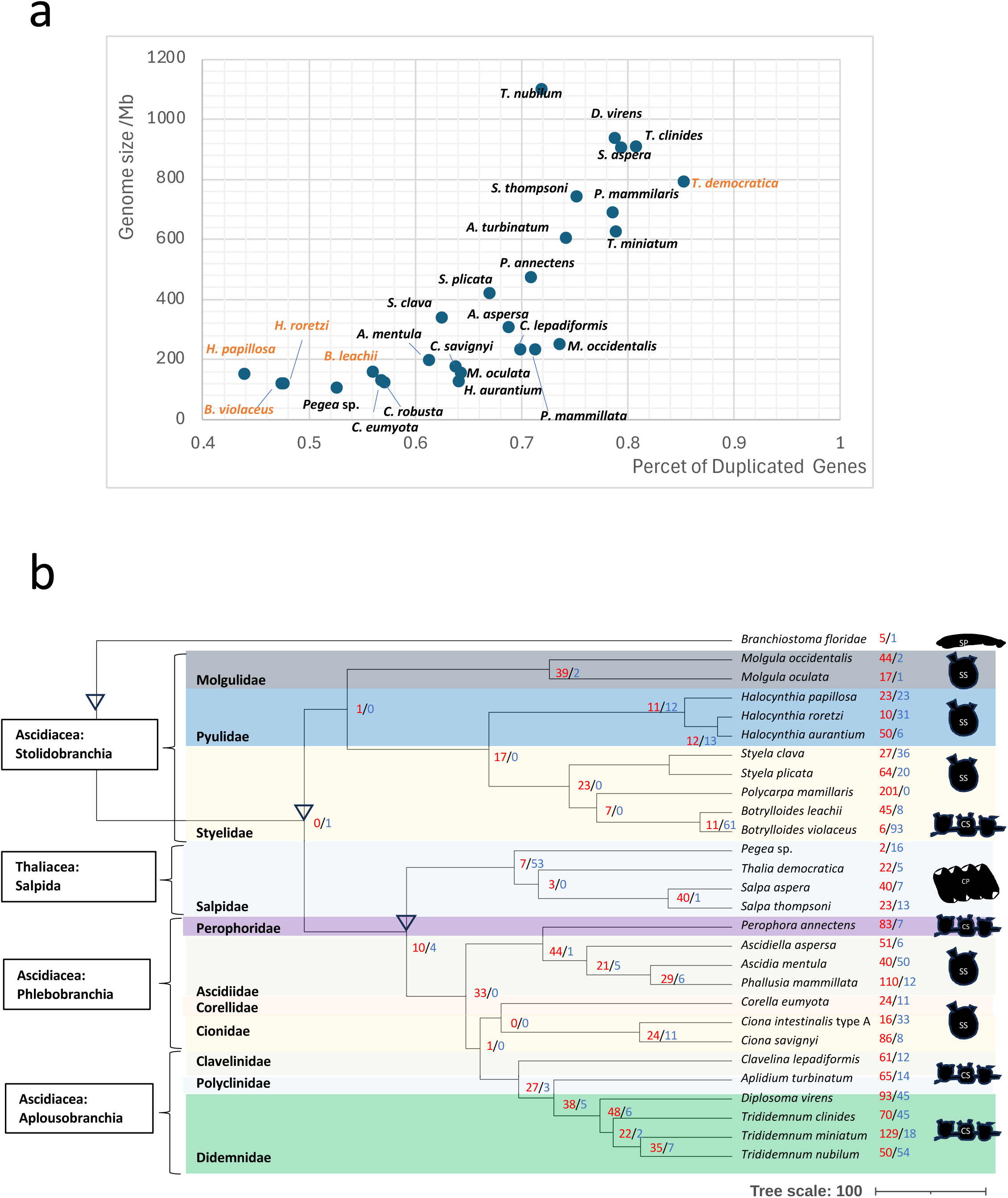
Phylogeny of tunicates and duplicated gene families. a) Scatter plot of duplicated gene ratio versus genome size in 28 tunicate species with gene models showing about 90% or higher BUSCO completeness. b) Phylogenetic tree constructed using the ML method with 27 tunicates and *Branchiostoma floridae* as the outgroup. Families are separated by different colors, and body plan or life cycles (SS: Solitary Sessile, CS: Colonial Sessile, SP: Solitary Planktonic, CP: Colonial Planktonic) of clades are illustrated at right. Time Calibration points are indicated by arrowheads. The numbers of highly expanded or contracted gene families are shown at each clade in red or blue.

### Phylogenetic analysis

We first conducted ortholog analysis using 39 species (38 tunicate species and *Branchiostoma floridae* as an outgroup) with OrthoFinder. Of the total 1,053,122 genes, 959,536 genes were assigned to 60,568 orthogroups. Because the 39 species did not share any single copy orthologs, shared 1,518 orthogroups containing paralogs derived from all 39 species were used to construct the species tree using STAG (Fig. S3a). The resulting topology supported recent class-level analyses such as the paraphyly in Ascidiacea, including Thaliacea [1]. The monophyly of Appendicularia and its internal relationships were also supported in recent phylogenetic studies [70] and the monophyly of Phlebobranchia and Aplousobranchia was consistent with prior work [71], while STAG analysis contains the effect of paralogs.

To minimize the impact of paralogs, species were ranked by the BUSCO completeness of their gene models, and those with lower scores were progressively excluded to maximize the number of single-copy orthologs (Table 8). We reduced the number of species from 39 to 32 (BUSCO > around 80%), 30 (BUSCO > 85%), 29, and 28 (BUSCO > around 90%), which yielded 44,324, 42,405, 42,405, and 39,311 orthogroups, respectively (Table 8). The number of single-copy orthogroups progressively increased to 94, 217, 240, and 385 as the number of species decreased.

Consequently, 28 tunicate species were chosen, and 385 single-copy orthologs were used for phylogenetic analysis based on maximum likelihood analysis using IQ-TREE (Fig. 1b; Fig. S3b). Bootstrap values supported the tree with nearly 100% except for the nodes of Corellidae and Cionidae, which had 94% support (Fig. S3b). ML Analysis with other species subsets (Table S8) produced the same topology.

In the resulting tree, Stolidobranchia was the earliest branching group, followed by Thaliacea, then the clade comprising Ascidiidae and Perophoridae, and finally the group of Corellidae, Cionidae, and Aplousobranchia. This tree suggests that Aplousobranchia was nested within the paraphyletic Phlebobranchia, which is inconsistent with STAG phylogeny (Fig. S4). This paraphyly of Phlebobranchia is consistent with recent studies although the relationships of families differ [1, 47, 72] (Fig. S4). Delsuc et al. [1] and Shenkar et al. [72] positioned Ascidiidae as the sister group of Cionidae and the Aplousobranchia clade, consistent with our tree, although [72] placed Corellidae as a sister group of Ascidiidae in contrast to our tree, which suggests the sisterhood of Corellidae and Cionidae. On the other hand, prior work [47] placed Cionidae as a sister group of Aplousobranchia, Ascidiidae, and Corellidae clade, while others [71] positioned Aplousobranchia as a sister group of Ascidiidae, Corellidae, and Cionidae (Fig. S4). These four studies did not include Perophoridae, and our study suggests a sister group relationship between Ascidiidae and Perophoridae.

### Gene duplication analysis

The percentage of duplicated genes was calculated by the results from OrthoFinder and plotted against genome size (Fig. 1a). Higher genome sizes corresponded to higher numbers of duplicated genes. Two species in *Halocynthia* and two species in *Botrylloides* exhibited low duplication rate (44%–56%). In contrast, *T. democratica* showed an approximately 85% duplication rate. These tendencies were preserved in the scatter plot of duplication rate and total gene number (Fig. 1a).

The numbers of rapidly expanded or contracted gene families were investigated by CAFÉ and mapped onto the phylogenetic tree of the 28 species (Fig. 1b). Interestingly, most gene families showed rapid expansion rather than contraction. Exceptions were observed in *Botrylloides* and Thaliacea, which exhibited rapid contraction. *Botrylloides* species also show a low duplication rate in overall gene sets (Fig. 1a). By contrast, *Halocynthia* species, despite their low duplication rates, did not exhibit substantial gene family contraction. Gene ontology enrichment analysis suggested that rapidly contracted orthogroups in *Botrylloides* and Thaliacea contain some groups related to development, morphogenesis, and metabolism (Table S9).

Rapidly expanded gene families in *A. aspersa* and gene families unique to this species were also investigated. A total of 51 gene families were identified as rapidly expanded—of these, 21 showed significant similarity to human genes (BLASTP, e-value < 0.01). These families are mainly associated with DNA binding, hormone biosynthetic process, transcriptional regulation, and extracellular matrix organization (Table S9). In addition, 315 gene families were found to be unique to *A. aspersa*. Gene ontology analysis indicated that these genes are enriched in functions related to the extracellular matrix, endoplasmic reticulum, and the omega-hydroxylase P450 pathway.

### Comparison based on Genome size and GC content

Genome size and GC content from 38 tunicate species genomes were mapped first (Fig. 2a–b). Genome size varied 17-fold and ranged from 64.3 Mb in *Oikopleura dioica* to 1,100 Mb in *Trididemnum nubilum*. In Appendicularia, *Oikopleura vanhoeffeni* and *Mesochordaeus erythrocephalus* show relatively high genome sizes (643.7 Mb and 874 Mb, respectively). Thaliacean species, including *Thalia democratica* (790.7 Mb), *Salpa aspera* (903.3 Mb), and *Salpa thompsoni* (742.8 Mb), also exhibit large genome sizes. *Pegea* sp. is an exception with a small genome size of 105.8 Mb. Among ascidians, seven of the nine species with genomes larger than 400 Mb were colonial.

**Fig. 2.**
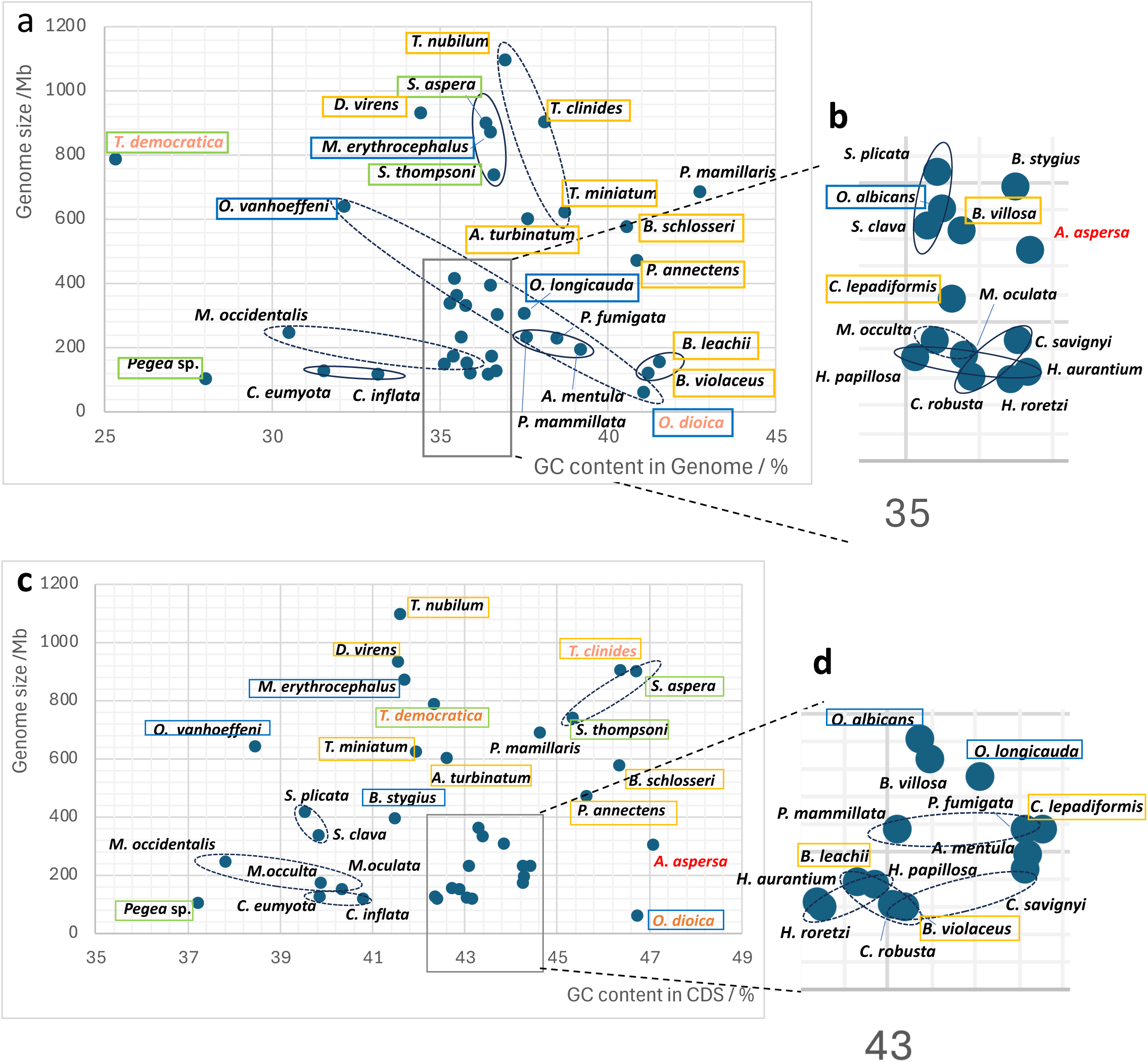
Comparison of genome size and GC contents. a) Scatter plot of genome size versus GC content in whole genomes among all 38 tunicate species. Species in the same genus were clustered. b) Close-up view of the crowded area within the orange rectangle in a). c) Scatter plot of genome size versus GC content in coding regions (CDS). d) Close-up view of the crowded area within the gray rectangle in c). Species positions that drastically changed are exhibited as red or orange colors. Species names are enclosed in rectangles indicating their lifestyles: Solitary sessile, no rectangles; Colonial sessile, yellow; Solitary planktonic, blue; Colonial planktonic: green.

Genome GC content varied from 25.3% in *T. democratica* to 42.7% in *Polycarpa mamillaris.* Thaliacean species, including *T. democratica* and *Pegea* sp., show very low GC content values (25.3% and 28.0%). In contrast, six species (*P. mamillaris*, *Botrylloides violaceus*, *Botrylloides leachii*, *Oikopleura dioica*, *Perophora annectens*, and *Botryllus schlosseri*) exhibited relatively high GC content above 40%—four of these belong to the family Styelidae.

A dense cluster of species was observed in the region of genome sizes between 120 Mb and 400 Mb and GC content between 35% and 37% (Fig. 2b). This cluster included two appendicularians (*Oikopleura* albicans and *Bathochordaeus stygius*) and 11 ascidian species. Of these 11 ascidians, all were solitary except for the colonial *Clavelina lepadiformis*. Genera represented by more than two species generally clustered together (Fig. 2a–b). Species within the same genus were mapped relatively close to each other. Notable exceptions were *Molgula occidentalis*, *Oikopleura* species, and *Trididemnum* species, which were dispersed among genera; three species in the same genus (*Trididemnum* and *Oikopleura*) were arranged linearly. In general, genus-level clusters exhibited a downward-right slope except in closely related genera such as *Ciona*, *Styela*, and *Botrylloides*.

GC content in coding regions (CDS) shows different trends compared to whole-genome GC content. GC content in CDS ranged from 37% to 47% typically 5% to 10% higher than genomic GC within the same species (e.g., 28.0% to 37.2% in *Pegea* sp.) (Fig. 2c–d; Table S9). While the relative positions among species were mostly conserved, several species (*T. democratica*, *T. clinides*, *A. aspersa*, *O. dioica*) exhibited marked shifts. Despite its extremely low genomic GC content, CDS GC content in *T. democratica* showed an intermediate value with an increase of almost 17% (Fig. 2c). *T. clinides* and *O. dioica* also shifted in relative position within their genera. Interestingly, *A. aspersa* exhibited the highest CDS GC content among all 38 genomes (47.1%) whereas its genomic GC content was intermediate (Fig. 2).

### GC content of tunicate genomes in different codon positions

To further investigate GC variation, we calculated GC content at the first, second, and third codon positions (GC1, GC2, GC3, respectively) for each species (Table S9). On average, GC1 showed a relatively high value of 48.3% while GC2 was relatively low at 38% (Fig. 3a). Interestingly, compared to GC1 and GC2, GC3 exhibited a wide distribution ranging from 29.9% (*Pegea* sp.) to 50.2% (*O. dioica*). GC contents across all regions were positively correlated, especially GC3, which showed the strongest correlation with CDS GC content (correlation coefficient = 0.975; Fig. 3b). Kernel density estimation of genome GC content revealed a major peak around 35% in both Appendicularia and Ascidiacea. In these two classes, GC peaks were also evident in CDS and GC3, whereas Appendicularia displayed a bimodal peak in GC1 and a downward shift in GC2. In contrast, Thaliaceans exhibited different positions—particularly with low values in *Pegea* sp. (Fig. 3b).

**Fig. 3.**
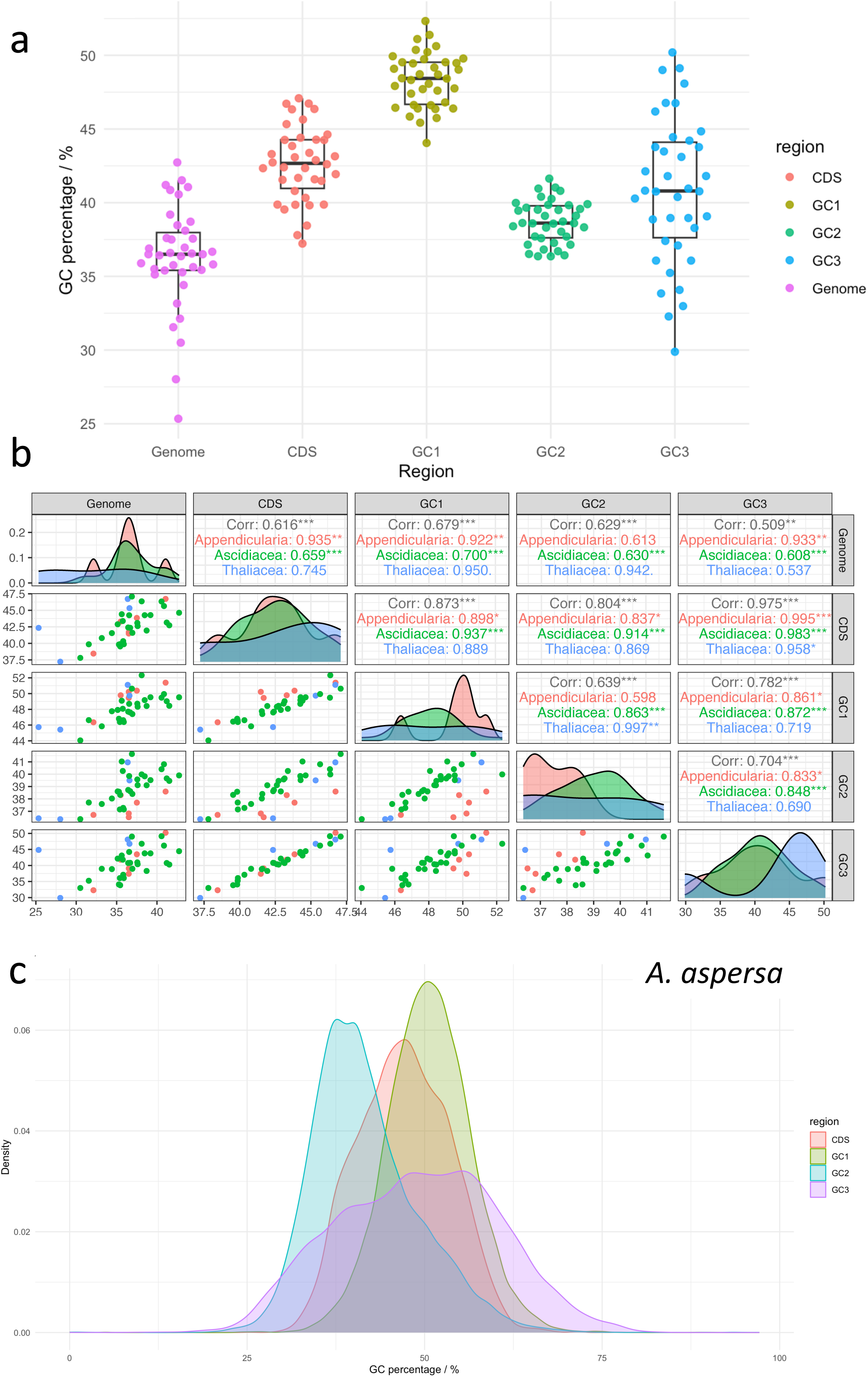
GC content in different positions of codons. a) Box plot of GC content in different regions of genomes. GC1–3 refers to GC content at the first, second, and third positions of codons. One dot represents the average value of one species. b) Scatter plot, density plot, and correlations of each combination of GC contents in different genome regions. Red, green, and blue colors correspond to Appendicularia, Ascidiacea, and Thaliacea, respectively. c) Density plot of GC content in all genes from different positions in the *A. aspersa* gene model (KAS25 model).

We next examined for GC content distributions for each gene in individual species (Fig. S5). In most species, GC1–3 density profiles formed sharp, unimodal peaks, while *O. longicauda*, *T. clinides*, *C. lepadiformis*, and *C. eumyota* displayed bimodal distributions. *O. longicauda* exhibited two peaks at around 30% and 52%, while *C. eumyota* showed a small second peak at about 50% across GC1–3. *T. clinides* displayed a second peak at about 60% only in GC1 and GC3. *C. lepadiformis* exhibited multiple minor peaks on the upper side of the main peak. Among Appendicularia, *O. dioica* showed a GC3 peak shift upwards. In Thaliacea, GC3 in *T. democratica* displayed a relatively low, rounded peak shape. In Phlebobranchia, *A. aspersa* showed a gradually shifted peak compared to other species (Fig. 3c; Fig. S5d). In Stolidobranchia, the shapes of the curves were almost identical except for a lower shift in GC3 of *M. occidentalis*.

Overall, GC1 tended to be relatively high, GC2 relatively low, and GC3 showed the widest variation among species. Notably, GC3 most strongly reflected interspecific differences and correlated closely with the overall CDS GC content.

### Codon usage bias in tunicate genomes

Codon usages of all 38 tunicates were calculated (Fig. S6). By combining usage fractions in all codons, we performed principal component analysis (PCA) and extracted PC1 and PC2 (Fig. 4a). These components accounted for 58.8% and 13.3% of the overall variance, respectively. Appendicularians were distant from other tunicates except for *O. vanhoeffeni* and *O. dioica*. Similarly, Pegea was positioned separately from other Salpidae species. Three *Trididemnum* species in Didemnidae, three molgulidae species, and four Pyulidae species exhibited clustering in the PCA map (Fig. 4a; Table S11). Interestingly, *A. aspersa* was isolated from other ascidiids (Fig. 4a).

**Fig. 4.**
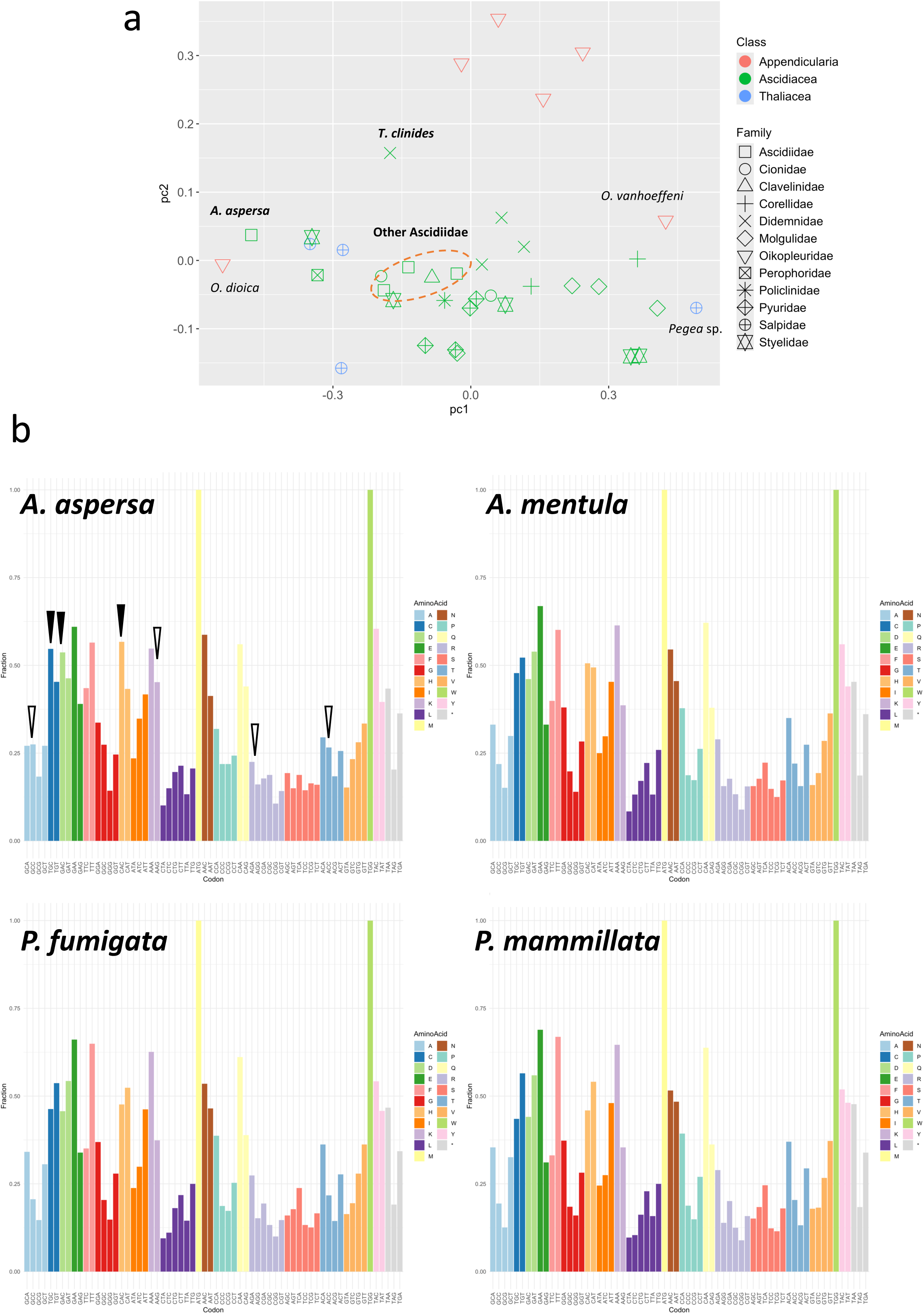
Codon usage analysis in tunicate species. a) Principal component analysis was performed on codon usage data of 38 tunicate species. Red, green, and blue colors correspond to Appendicularia, Ascidiacea, and Thaliacea, respectively. Different shapes of dots correspond to all families of tunicates. PC1 and PC2 accounted for 58.8% and 13.3% of the overall variance, respectively. b) Codon usages of amino acids in four ascidiids. The vertical axis show the fractions of each codon. Black pins indicate the reversal of codon usage compared to other Ascididae species. White pins indicate codon usage changes that are not as significant as reversals.

Versus other Ascidiidae, *A. aspersa* exhibited some changes in codon usage fractions (Fig. 4b). For example, in codons for cysteine (C), *A. aspersa* prefers TGC than TGT while other Ascidiidae species favor TGT. Similarly, *A. aspersa* prefers GAC rather than GAT for aspartic acid (D), while other species show the opposite tendency. Additionally, *A. aspersa* favours CAC was preferred than CAT for histidine (H) codons. Fractional changes were also notable in alanine (A), lysine (K), arginine (R), and threonine (T). Many of these changes involved a conversion from A/T to G/C at the third codon. These tendency switches were also observed relative to other phlebobranchs.

Codon usage variation analysis in related species is also remarkable in *O. dioica,* contrary to other Appendicularia species (Fig. S6a). Tendency changes were observed in cysteine (C), phenylalanine (F), histidine (H), isoleucine (I), asparagine (N), and tyrosine (Y), sharing C and H with *A. aspersa.* In the AT-rich *Pegea* sp., significant switches in usage tendency were only observed in asparagine (N), but the usage bias was enhanced in many codons (Fig. S6b). In Phlebobranchia, Corella species shared codon bias switches compared to other phlebobranchs particularly in asparagine (N). Aplousobranchia, Stolidobranchia species exhibited fewer changes in codon usage bias.

ENC-GC3 plots were drawn for codon usage variation analysis (Fig. 5; Fig. S4). A larger ENC value indicates a stronger bias in codon usage and a smaller ENC value indicates equal usage among codons. Mutational effects may be the dominant factor in the codon preference of genes on the expected curve [66]. A position below the curve indicates that the preference is under selection pressure. Distributions of genes in most Appendicularia is comparatively below the left of the peak (Fig. S7a). In contrast, *O. dioica* shows a clear tendency of extending down to the right indicating some popularity of genes under the influence of natural selection. *O. longicauda* shows high GC genes but is relatively positioned higher compared to *O. dioica*, which may be driven by sequence.

**Fig. 5.**
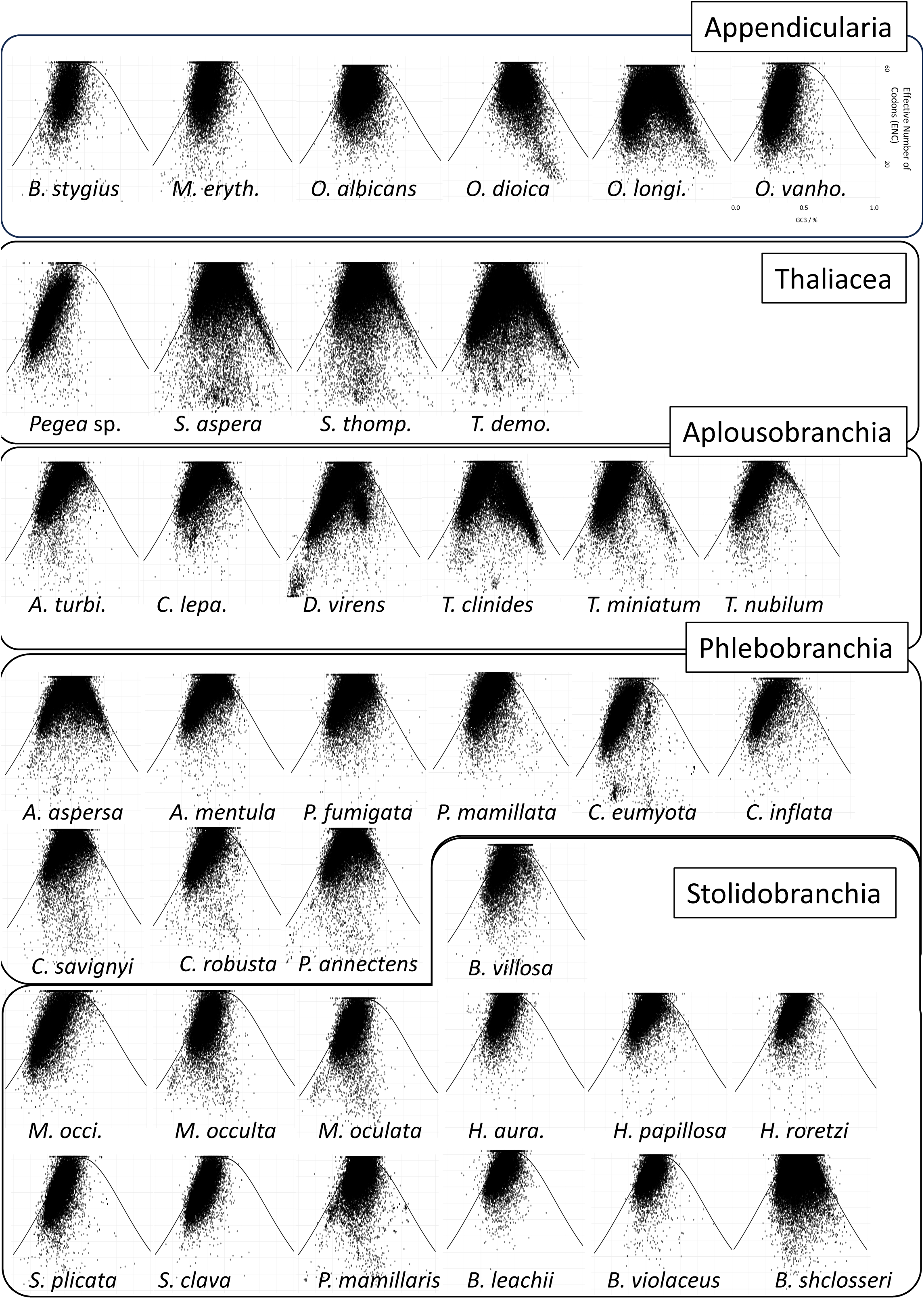
ENC-GC3 plot of 38 tunicates. Scatter plot of effective codon numbers (ENC) and GC content in the third codon position (GC3) in 38 tunicates. Axes were omitted almost and the example in *Oikopleura vanhoeffeni* is shown (Right Upper). One dot corresponds to one gene. The theoretical line was plotted according to the description in the methods section.

Among Thaliacea, *Pegea* sp. is distinctly positioned on the left side of the standard curve (Fig. S7b). Plots of the other three thaliaceans had similar shapes extending to the right side along the standard curve. Aplousobranchia species show almost left-side distributions along the curve, but *T. clinides* displays a right distribution similar to the left. The tendency of right distribution is weakly observed in other *Trididemnum* species, *T. miniatum*, and *T. nubilum* (Fig. S7c).

Phlebobranchians show distributions at the top of the peak and extending left except for *Ascidiella aspersa*, which has a right distribution (Fig. S7d). Stolidobranchians show almost AT-side distributions along the standard line distribution except for *B. schlosseri* (Fig. S7e).

Overall, codon usage in ascidians reflects patterns characteristic of each family with some exceptions, such as *A. aspersa*, *O. dioica,* and *Pegea* sp., which show clear preferences diverging from closely related species (Fig. 4; Fig. 5).

### Web-based database enables genome-wide identification and transcriptomic investigation

We integrated all genome sequences, and then constructed gene models, functional annotations, and ortholog analyses of all 38 tunicate species into a new tunicate genome database named as TUNOME (https://ciona.bpni.bio.keio.ac.jp/Tunome/Latest/). Functional annotations can be easily searched using various queries, such as gene ID, common names, or domain names (Fig. 6b; Fig. S8a). Corresponding to a selected gene, expression levels were represented with transcripts per million (TPM) of *A. aspersa* are graphically showed, and both nucleotide and protein sequences are available for download (Fig. 6c; Fig. S8b). A genome browser is provided and enables visualization of RNA-seq alignments derived from different organs or embryonic stages (Fig. 6d; Fig. S8c). The phylogram and all relative sequences of the orthogroup containing the selected gene can also be viewed (Fig. 6e; Fig. S8d). Additionally, BLAST searches can be performed against all 38 genomes and their gene models (Fig. 6f; Fig. S8e). The “Blast to all species” mode allows simultaneous searches across all species, and both summary tables and hit sequences can be downloaded (Fig. S8f). Initially, we tried “Blast to all species” mode with Blastp to search DNA repair genes into 38 tunicate genomes. Using Rscipt enables in download page, we visualized possession of genes involved in each pathway (Fig. S9). In terms of Base Excision Repair, specific gene lacks were observed in Larvacea, Thaliacea, and Styelidae (Fig. 6g). These gene lacks were also confirmed by tblastx to genomes (Fig. S9f).

**Fig. 6.**
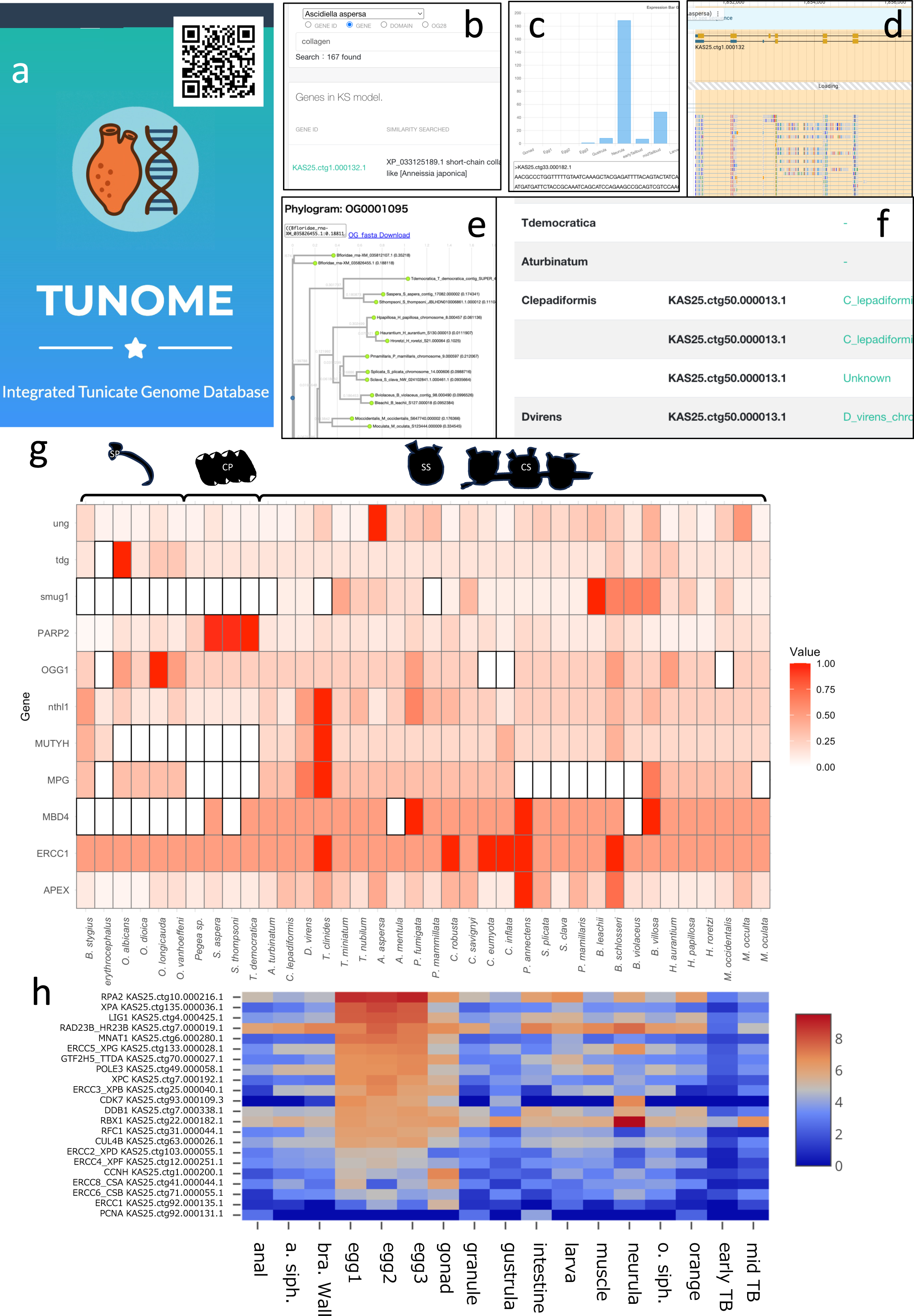
TUNOME database and application for DNA repair genes. a) TOP page of TUNOME database, suitable for PC and smartphone. b) annotations page, which provide searching for annotations of all gene models from 38 tunicate genomes. c) genome browser page linked to each gene in a). d) expression bar graph displayed in a). e) phylograms linked to each gene in a). f) Blast result page, which provide blast to 38 tunicates at once. g) DNA repair gene sets related to Base Excision Repair (BET) in all tunicates obtained using blastp to all function and R script downloaded from Download page. Color show relative value of possession of each gene. White squares with black outline show correspond species doesn’t have correspond gene in constructed gene model. h) Heatmap of NER genes expression in *A. aspersa* visualized in “expression data” page in TUNOME. Color values indicate log2 folded TPM. Genes were sorted by the order of the average values of TPM in three eggs samples.

We also developed “ortholog search” mode in TUNOME based on OrthoFinder analysis to narrow down some clade specific genes (Fig. S10a). We first gained 145 orthogroups conserved only in ascidiid genomes, and obtained total of 259 *A. aspersa* genes belonging to the 145 orthogroups by “convert OG names to gene names” page (Fig. S10b). Using the “Expression Data” page, 259 genes were sorted by the order of abundance in expression in *A. aspersa* eggs and visualized as a heatmap (Fig. S10c–d). There were 30 TOP genes, and 19 genes have no annotations (Table S13). Blastp search of 38 species was done using TUNOME and showed two genes (KAS25.ctg15.000028.1, KAS25.ctg85.000038.1) were confirmed to be conserved only in four ascidiids. These have eggs or embryo specific expression in *A. aspersa* (Fig. S10e–g).

We also explored transcripts abundant in *A. aspersa* eggs using “expression data” page (Fig. S11a–b). In 30 genes TOP rank of transcripts, almost genes related to before or after fertilization process such as cell cycle regulation (CCNA2, CCNB2, CDC6, CCNE1, ERH, 14-3-3, ZAR1L), GPCR/Calcium pathway (CALM1, CAM, GNAT3), cell adhesion (CLDN1, CTNNB1), energy metabolism in mitochondria (SLC25A4, VDAC1, CYCS), RNA homeostasis (DDX5, DDX39, ALYREF), translation regulation (ZAR1L, EEF1A1), spindle assembly (TUBA1B). Also, the ubiquitine proteasome pathway (FBXL12, SPSB1), dNTP synthesis (RRM2B), transmembrane trafficking protein (TMED10), glutamine synthetase (GLUL), and chaperone (P4HB), systems were seen besides there are uncharacterized transcripts highly abundant in eggs (KAS25.ctg60.000084.1, KAS25.ctg39.000122.1, KAS25.ctg3.000148.1) (Table S14). We also investigated expression levels of nucleotide excision repair (NER) pathway genes and found that most of them are rich in *A. aspersa* eggs (Table S15; Fig. 6h).

## Discussion

### Improving construction of gene models from GC rich tunicate genomes

Gene prediction could be challenging due to high GC content, as observed in the gene prediction of the western honey bee genome with bimodal GC content [73]. *T. clinides* also exhibits bimodal GC content, and several other species representing *A. aspersa*, have relatively high GC values in their CDS. In our analysis, *T. clinides* and *A. aspersa* have shown relatively lower BUSCO value in gene models constructed by BRAKER2, but it was suitable for various species including *A. mentula*, *P. mammillata*, and *B. violaceus,* which generated gene models that included nearly all BUSCO genes predicted across the whole genome (Table S12). In these cases, GeMoMa performed well producing BUSCO scores around 90% (Table S12). Our run in GeMoMa ignored missing start/stop codons, which almost explained the gaps observed in some species (*S. thompsoni*, *S. aspera*, and *P. annectens*) (Table S12), while there are still gaps between BRAKER2 and GeMoMa in *T. clinides* and *A. aspersa*.

### Variation in genome size and gene numbers in tunicates

Consistent with our results, some reports indicate a positive relationship between genome size and predicted gene size although protein-coding regions contribute only a small fraction to overall genome size [74]. Genome size and organism complexity are not necessarily related, but cell size and cell division rate are associated; larger genome sizes are observed in the nuclei of larger and slowly dividing cells [74]. Compound species *Botryllus schlosseri*, which has a relatively large genome in tunicates (about 580 Mb), exhibits slow embryo development compared to *Ciona robusta*, which has a smaller genome (about 123 Mb) [75]. To verify the existence of a constraint on developmental speed and genome size, further investigation of embryogenesis in other compound ascidian species with larger genome sizes is required, as this remains unclear in most species.

### The phylogeny of the Tunicata

Phylogenetic analysis in tunicates has been complicated by accelerated rates of evolution in their nuclear and mitochondrial genomes, which could result in long-branch attraction (LBA) [1]. The long branch observed in *Oikopleura dioica* reflects the high evolutionary rate of this species complicating the debate regarding its basal position as the sister group of all other tunicates [76]. Additionally, 18S rRNA sequences of Aplousobranchia species seem to evolve at high rates causing long branches [77]. In our STAG tree, long branches were not observed in Appendicularians suggesting that transcriptomic-level analysis might resolve the long branch effects in tunicates, although this analysis contains the effects of paralogs (Fig. S3a). Long branches were not seen in Aplousobranchia in the ML tree, and Thaliacea contained a slightly long branch (Fig. S3b).

Two main unresolved phylogenetic questions in tunicates are: 1) the relationships among the three classes of tunicates, and 2) the relationships between Phlebobranchia and Aplousobranchia. For the first question, reliable appendicularian genomes are necessary because there are still no reliable phylogenetic analysis based on ML or Bayesian. Salpidae in Thaliacea were placed within Ascidiacea with 100% bootstrap support in our analysis (Fig. 1b; Fig. S3b) consistent with recent literature, while the monophyly of Thaliacea should be proven through phylogenomics approaches. Regarding the second question, there is no consensus on the relationships among families in Phlebobranchia and Aplousobranchia with completely reliable bootstrap values (Fig. S4). To address this problem, comprehensive sampling of various taxa is required because there are few representatives from five families in Phlebobranchia, including Agneziidae and Plurelidae—many of which inhabit deep-sea environment. The problem is even more serious in Aplousobranchia, which has the highest number of species and families in tunicates, making species identification relatively difficult [78]. For phylogenomic approaches, several alternative strategies may be helpful: using genes that are not necessarily single-copy [79] and accurately identifying orthologs [80].

### Variety of GC Content in tunicate genomes

A study reported a mean GC content of 39.9% for sampled metazoan genomes, although some species show unusually high or low GC content such as the hydroid *Hydra magnipapillata* (29% GC) and the sea lamprey *Petromyzon marinus* (46% GC) [81]. Interestingly, tunicates exhibit a wide range of genomic GC content within a single subphylum ranging from 25.3% in *T. democratica* to 42.7% in *P. mamillaris.* In the scatter plot of genome size versus genomic GC content, species within the same genus were either clustered closely or dispersed along a downward-sloping trend line in genera with more than two species such as *Trididemnum* and *Oikopleura* (Fig. 2a). A negative correlation between genome size and GC content has also been reported in monocot genomes possibly due to the higher energetic cost of nucleotide synthesis, which causes a deficiency of dGTP and dCTP, leading to the misincorporation of the less costly dATP and dTTP, and an overall mutation bias toward AT-rich DNA [82].

Regarding coding regions, the GC content was generally more than 5% higher than that of the whole genome ranging from 29.9% to 50.2% (Fig. 2c–d). These values of CDS GC content were relatively low compared to a dataset of 33 mammals, which ranged from 43.9% to 57.9% [83]. Tunicate genomes exhibited higher GC content at the first codon position (GC1) than at the second codon position (GC2); while the third codon position (GC3) showed a wider range among tunicate species compared to GC1 and GC2 (Fig. 3a). GC indices have positive correlations with each other, especially GC3, which has a high correlation with CDS GC content (Fig. 3b). This tendency is also observed in the prokaryotes and viral genes [84].

Beyond genus level conservation in genome and GC relationships, there were also class-level tendencies in GC contents in Tunicata species. Appendicularia and Ascidiacea shares GC peak in genome, CDS, and GC3 regions, but this is not seen GC1 and GC2 (Fig. 3b). Thaliacea have shown more uniformly spread line, but an obviously higher GC3 peak compared to Appendicularia and Ascidiacea. These tendencies reflect distinctive genomic evolutionary traits they experienced, which contains different features of Thaliacea and could raise suspicion on the position of one monophyly in Ascidiacea.

### Biological significance of GC Content

The biological significance of GC variation in tunicates remains unresolved. However, our comparative analyses showed that GC-related genomic features differ systematically among the major tunicate classes rather than varying randomly among species (Figs. 2–5). Ascidiaceans generally exhibited moderate genomic GC content, and both Ascidiacea and Appendicularia showed major peaks in genome GC, CDS GC, and GC3. Appendicularia also displayed distinctive features, including a bimodal GC1 distribution and a downward shift in GC2. Thaliaceans showed a different overall distribution from those of Ascidiacea and Appendicularia, including low genomic GC values in some species. These results suggest that GC content, GC3, and codon usage reflect class-associated modes of genome evolution across tunicates [81].

One implication of this pattern is that class-level differences in nucleotide composition are accompanied by differences in coding-sequence properties. Because GC3 is closely linked to synonymous codon usage, the observed differences among Ascidiacea, Appendicularia, and Thaliacea are likely to affect codon preference and thus influence how coding information is structured in each lineage[85, 86]. In this sense, GC-related variation is not only a compositional property of genomes but also a potential indicator of lineage-specific coding constraints.

Our results also suggest that the mechanisms shaping GC bias may differ among tunicate classes. The class-associated patterns observed in genomic GC content, GC3, codon usage, and ENC-GC3 distributions are consistent with differences in the relative contributions of mutational bias, selection on codon usage, and GC-biased gene conversion (gBGC) [81, 86, 87]. However, the present study does not allow these mechanisms to be distinguished conclusively. Likewise, although these GC-related features may ultimately be linked to ecological or life-history traits, our current data support this only at the level of comparative genomic tendencies, not direct functional causation.

Overall, the present analyses show that GC content, GC3, and codon usage are informative comparative features for understanding genome evolution at the class level in tunicates. The shared tendencies identified within Ascidiacea, Appendicularia, and Thaliacea provide a useful framework for future work on the molecular mechanisms and biological consequences of nucleotide composition bias in this subphylum [81, 83, 86].

### DNA repairment

*O. dioica* has extreme genome scrambling caused by thousands of chromosomal rearrangements, which may be promoted by the loss of the canonical nonhomologous end-joining (NHEJ) DNA repair pathway among appendicularians [70, 88]. Our dataset also suggests a lack of DNA repair pathways containing NHEJ in appendicularians (Fig. S9). Interestingly, thaliacean species also share a lack of genes associated with DNA repair (smug1, MUTYH, MPG, BRCA1, XRCC4), although Appendicularia and Thaliacea are not sister groups. As suggested in appendicularians, potential losses in DNA repair pathways have occurred in the common ancestor of thaliaceans, possibly affecting the high variability in thaliacean genomes, which is represented by genome size, GC value, and the rapid evolutionary rate observed in the maximum likelihood tree (Fig. S3b). Chromosomal rearrangement can be caused in Thaliacean, which cannot be verified because there are limited chromosomal data of these species. Base excision repair (BER), which includes smug1, MUTYH, and MPG, is thought to increase GC to AT transitions when impaired [89, 90]. AT-rich genomes in *Pegea* sp. could be caused by a lack of BER pathway as seen in the deep-sea bone-eating polychaete worm that has AT-rich genome and an underlying loss of genes encoding components of the BER pathway represented by SMUG1 [81, 89].

### The genomic and transcriptomic basis enables further insight into tunicate characteristics

We constructed a genome assembly of *A. aspersa* and gene models across 38 tunicates. Using our resources integrated to TUMOME database made comparing tunicate species genomes including non-model animals easy (Fig. S6g). We also combined ortholog analysis and transcriptome into TUNOME database, enabling bridge comparative eco-evo-devo: These resources can also be helpful for developmental experiments. Furthermore, orthogroup analyses anchored by our gene models can aid investigations into shared features, such as bio-transparency in ascidian embryos.

To demonstrate this, we investigated ascidiids-specified orthogroups using OrthoFinder to study mechanisms of eggs transparency of Ascidiidae species. This led to 145 orthogroups, corresponding to 259 genes of *A. aspersa* with 19 unannotated genes highly expressed in eggs (Table S13). On the other hands, blastp confirmation showed the possibility of existing other related genes in outer ascidiids; these were not discovered at orthofinder analysis (Fig. S10e). In the future, more concise identification of orthologs should be done with another methods [80], although orthofinder analysis has a great merits in terms of easy large-scale comparison in easy way.

In *A. aspersa* eggs, mRNA can be translated in unfertilized period, which could be related to maintaining the eggs’ remarkable transparency [9, 91, 92]. We obtained TOP 30 genes that were highly abundant in *A. aspersa* eggs (Fig. S11a–b; Table S14). Most of them are thought to be related to fertilization process (cell cycle regulation, GPCR/calcium, energy metabolism, and spindle assembly) although transcription before fertilization possibly enables maintaining eggs transparency. Also, nucleotide excision repair (NER) genes were highly expressed especially in eggs of *A. aspersa* (Fig. 6h). NER pathway plays an important role in DNA repair induced by UV radiation [93], which can protect eggs from UV damage to transparent eggs which is potentially vulnerable to UV. More application of our database will facilitate experiments in non-model animals and enable species comparisons among tunicate traits.

### Conclusions

We constructed genomic and transcriptomic resources for *A. aspersa*. Other tunicate species’ genomes were also annotated, and gene models were rebuilt achieving high-quality gene models containing 27 species with BUSCO scores greater than 80%. Phylogenetic analysis produced a new hypothesis in tunicate phylogeny, and gene duplication analysis characterized multiple taxa within tunicates. Genome statistics of tunicates exhibited high variability in genome size and GC content including high GC content in the CDS of *A. aspersa* genome and codon usage bias. We constructed TUNOME database to easily compare tunicate species genomes. Using this, we discovered DNA repair pathway lack in Thaliacea and potentially candidate genes related to eggs transparency mechanisms in Ascidiidae. These results can accelerate evo-devo research through species comparisons including non-model organisms.

### Data availability

Genome sequencing data and assembly were deposited to BioProject accession PRJNA1345923. Gene models and protein sequences for other species were deposited in the TUNOME database: https://ciona.bpni.bio.keio.ac.jp/Tunome/Latest/

## Author Contributions

KH conceived the project. KH and TS designed the experiments. TS and SK performed experiments with living samples. KN, YN performed the sequencing. VJ and YS conducted the assembly. TS, SK, SM, and KN carried out the gene model construction. TS performed subsequent analysis and constructed the website. TS and KH drafted the manuscript. KO, YS, KN, and KH critically revised the manuscript and supervised the research. All authors reviewed the manuscript.

## Acknowledgements

We thank Dr. Minoru Ikeda and Captain Toyokazu Hiratsuka (Onagawa Field Center of Tohoku University); Dr. Makoto Kanamori, Dr. Masafumi Natsuike, and Takuya Mizukami (Hokkaido Research Organization, Hakodate Fisheries Research Institute); Dr. Gaku Kumano (Graduate School of Life Sciences, Tohoku University); and Akio Takiya, Kazuya Sakai, Dr. Tatsunari Mori, Dr. Takaaki Kayaba, and Dr. Motohito Yamaguchi (Hokkaido Research Organization, Central Fisheries Research Institute) for their assistance in collecting samples. We also thank Dr. Kazuho Ikeo and Dr. Sonoko Kinjo (National Institute of Genetics) and Dr. Takashi Makino (Tohoku University) and Naohiro Hasegawa (Hiroshima Shudo University) and Euichi Hirose (University of Ryukyus) for their helpful comments. We thank Mr. Wu Jiayang for building the computer for the analysis. We thank Ms. Haruka M. Funakoshi and Ms. Yuki S. Kogure for handling *Ascidiella aspersa* samples. We are grateful to Dr. Ken Naito (The University of Tokyo), Dr. Taro Maeda (Keio University), and Dr. Jun Inoue (The University of Tokyo) for sharing valuable articles about genome analysis methods on their website. We would like to express our sincere gratitude to Dr. Nori Satoh for sequencing support.

## Abbreviations

GC1: GC content in the first codon
GC2: GC content in the second codon
GC3: GC content in the third codon
gBGC: GC-biased gene conversion
ENC: Effective number of codons
PCA: Principal component analysis
SS: Solitary Sessile
CS: Colonial Sessile
SP: Solitary Planktonic
CP: Colonial Planktonic

**Supplementary Material 1**

SupplementaryMaterial_1.xlsx. Table S1–S12.

**Supplementary Material 2**

SupplementaryMaterial_2.docx. Fig. S1–S7.

**Supplementary Material 3**

SupplementaryMaterial_3.docx. Fig. S8–S9.

## References

1. Delsuc F, Philippe H, Tsagkogeorga G, Simion P, Tilak M-K, Turon X, et al. A phylogenomic framework and timescale for comparative studies of tunicates. BMC Biol. 2018;16:39.

2. Satoh N. Developmental Genomics of Ascidians. Nashville, TN: John Wiley & Sons; 2014.

3. Dardaillon J, Dauga D, Simion P, Faure E, Onuma TA, DeBiasse MB, et al. ANISEED 2019: 4D exploration of genetic data for an extended range of tunicates. Nucleic Acids Res. 2020;48:D668–75.

4. Yasuo H, McDougall A. Practical Guide for Ascidian Microinjection: Phallusia mammillata. Adv Exp Med Biol. 2018;1029:15–24.

5. Shito TT, Hasegawa N, Oka K, Hotta K. Phylogenetic comparison of egg transparency in ascidians by hyperspectral imaging. Sci Rep. 2020;10:20829.

6. Lynch SA, Darmody G, O’Dwyer K, Gallagher MC, Nolan S, McAllen R, et al. Biology of the invasive ascidian Ascidiella aspersa in its native habitat: Reproductive patterns and parasite load. Estuar Coast Shelf Sci. 2016;181:249–55.

7. Palanisamy SK, Thomas OP, P McCormack G. Bio-invasive ascidians in Ireland: A threat for the shellfish industry but also a source of high added value products. Bioengineered. 2018;9:55–60.

8. Nishikawa T, Oohara I, Saitoh K, Shigenobu Y, Hasegawa N, Kanamori M, et al. Molecular and morphological discrimination between an invasive ascidian, Ascidiella aspersa, and its congener A. scabra (Urochordata: Ascidiacea). Zoolog Sci. 2014;31:180–5.

9. Funakoshi HM, Shito TT, Oka K, Hotta K. Developmental Table and Three-Dimensional Embryological Image Resource of the Ascidian Ascidiella aspersa. Frontiers in Cell and Developmental Biology. 2021;9. 10.3389/fcell.2021.789046.

10. Kogure YS, Hotta K. An improved fertilization protocol for*Ascidiella aspersa*: Promoting a new chordate model in developmental biology. bioRxiv. 2025;:2025.03.10.642315. 10.1101/2025.03.10.642315.

11. Dehal P, Satou Y, Campbell RK, Chapman J, Degnan B, De Tomaso A, et al. The draft genome of Ciona intestinalis: insights into chordate and vertebrate origins. Science. 2002;298:2157–67.

12. Ruan J, Li H. Fast and accurate long-read assembly with wtdbg2. Nat Methods. 2020;17:155–8.

13. Liu H, Wu S, Li A, Ruan J. SMARTdenovo: a de novo assembler using long noisy reads. Gigabyte. 2021;2021:1–9.

14. Kolmogorov M, Yuan J, Lin Y, Pevzner PA. Assembly of long, error-prone reads using repeat graphs. Nat Biotechnol. 2019;37:540–6.

15. Koren S, Walenz BP, Berlin K, Miller JR, Bergman NH, Phillippy AM. Canu: scalable and accurate long-read assembly via adaptive k-mer weighting and repeat separation. Genome Res. 2017;27:722–36.

16. Flynn JM, Hubley R, Goubert C, Rosen J, Clark AG, Feschotte C, et al. RepeatModeler2 for automated genomic discovery of transposable element families. Proc Natl Acad Sci U S A. 2020;117:9451–7.

17. Smit, AFA, Hubley, R & Green, P. RepeatMasker Open-4.0. 2013-2015.

18. Manni M, Berkeley MR, Seppey M, Simão FA, Zdobnov EM. BUSCO update: Novel and streamlined workflows along with broader and deeper phylogenetic coverage for scoring of eukaryotic, prokaryotic, and viral genomes. Mol Biol Evol. 2021;38:4647–54.

19. Nishimura O, Hara Y, Kuraku S. gVolante for standardizing completeness assessment of genome and transcriptome assemblies. Bioinformatics. 2017;33:3635–7.

20. Sensui N, Itoh Y, Okura N, Shiba K, Baba SA, Inaba K, et al. Spawning-Induced pH Increase Activates Sperm Attraction and Fertilization Abilities in Eggs of the Ascidian, Phallusia philippinensis and Ciona intestinalis. Int J Mol Sci. 2023;24:2666.

21. Andrews S. FastQC: a quality control tool for high throughput sequence data. 2010.

22. Shen W, Le S, Li Y, Hu F. SeqKit: A cross-platform and ultrafast toolkit for FASTA/Q file manipulation. PLoS One. 2016;11:e0163962.

23. Chen S, Zhou Y, Chen Y, Gu J. fastp: an ultra-fast all-in-one FASTQ preprocessor. Bioinformatics. 2018;34:i884–90.

24. Haas BJ, Papanicolaou A, Yassour M, Grabherr M, Blood PD, Bowden J, et al. De novo transcript sequence reconstruction from RNA-seq using the Trinity platform for reference generation and analysis. Nat Protoc. 2013;8:1494–512.

25. Kim D, Paggi JM, Park C, Bennett C, Salzberg SL. Graph-based genome alignment and genotyping with HISAT2 and HISAT-genotype. Nat Biotechnol. 2019;37:907–15.

26. Ito M, Ohashi H, Takemoto M, Muto C, Seiko T, Noda Y, et al. Single candidate gene for salt tolerance of Vigna nakashimae (Ohwi) Ohwi & Ohashi identified by QTL mapping, whole genome sequencing and triplicated RNA-seq analyses. Breed Sci. 2024;74:93–102.

27. Brůna T, Hoff KJ, Lomsadze A, Stanke M, Borodovsky M. BRAKER2: automatic eukaryotic genome annotation with GeneMark-EP+ and AUGUSTUS supported by a protein database. NAR Genom Bioinform. 2021;3:lqaa108.

28. Waterhouse RM, Tegenfeldt F, Li J, Zdobnov EM, Kriventseva EV. OrthoDB: a hierarchical catalog of animal, fungal and bacterial orthologs. Nucleic Acids Res. 2013;41 Database issue:D358–65.

29. Keilwagen J, Hartung F, Grau J. GeMoMa: Homology-Based Gene Prediction Utilizing Intron Position Conservation and RNA-seq Data. Methods Mol Biol. 2019;1962:161–77.

30. Pertea G, Pertea M. GFF Utilities: GffRead and GffCompare. F1000Res. 2020;9:304.

31. Hart AJ, Ginzburg S, Xu MS, Fisher CR, Rahmatpour N, Mitton JB, et al. EnTAP: Bringing faster and smarter functional annotation to non-model eukaryotic transcriptomes. Mol Ecol Resour. 2020;20:591–604.

32. Satou Y, Nakamura R, Yu D, Yoshida R, Hamada M, Fujie M, et al. A Nearly Complete Genome of Ciona intestinalis Type A (C. robusta) Reveals the Contribution of Inversion to Chromosomal Evolution in the Genus Ciona. Genome Biol Evol. 2019;11:3144–57.

33. Satou Y, Tokuoka M, Oda-Ishii I, Tokuhiro S, Ishida T, Liu B, et al. A Manually Curated Gene Model Set for an Ascidian, Ciona robusta (Ciona intestinalis Type A). Zoolog Sci. 2022;39:253–60.

34. Small KS, Brudno M, Hill MM, Sidow A. A haplome alignment and reference sequence of the highly polymorphic Ciona savignyi genome. Genome Biol. 2007;8:R41.

35. Voskoboynik A, Neff NF, Sahoo D, Newman AM, Pushkarev D, Koh W, et al. The genome sequence of the colonial chordate, Botryllus schlosseri. Elife. 2013;2:e00569.

36. Blanchoud S, Rutherford K, Zondag L, Gemmell NJ, Wilson MJ. De novo draft assembly of the Botrylloides leachii genome provides further insight into tunicate evolution. Sci Rep. 2018;8:5518.

37. Stolfi A, Lowe EK, Racioppi C, Ristoratore F, Titus Brown C, Swalla BJ, et al. Divergent mechanisms regulate conserved cardiopharyngeal development and gene expression in distantly related ascidians. Elife. 2014;3. 10.7554/eLife.03728.

38. Wang K, Tomura R, Chen W, Kiyooka M, Ishizaki H, Aizu T, et al. A genome database for a Japanese population of the larvacean Oikopleura dioica. Dev Growth Differ. 2020;62. 10.1111/dgd.12689.

39. Brozovic M, Dantec C, Dardaillon J, Dauga D, Faure E, Gineste M, et al. ANISEED 2017: extending the integrated ascidian database to the exploration and evolutionary comparison of genome-scale datasets. Nucleic Acids Res. 2018;46:D718–25.

40. Bliznina A, Masunaga A, Mansfield MJ, Tan Y, Liu AW, West C, et al. Telomere-to-telomere assembly of the genome of an individual Oikopleura dioica from Okinawa using Nanopore-based sequencing. BMC Genomics. 2021;22:222.

41. Naville M, Henriet S, Warren I, Sumic S, Reeve M, Volff J-N, et al. Massive changes of genome size driven by expansions of non-autonomous transposable elements. Curr Biol. 2019;29:1161–1168.e6.

42. Castellano KR, Batta-Lona P, Bucklin A, O’Neill RJ. Salpa genome and developmental transcriptome analyses reveal molecular flexibility enabling reproductive success in a rapidly changing environment. Sci Rep. 2023;13:21056.

43. Daric V, Lanoizelet M, Mayeur H, Leblond C, Darras S. Genomic Resources and Annotations for a Colonial Ascidian, the Light-Bulb Sea Squirt Clavelina lepadiformis. Genome Biol Evol. 2024;16. 10.1093/gbe/evae038.

44. Bishop J, Harley J, Mrowicki R, Marine Biological Association Genome Acquisition Lab, Darwin Tree of Life Barcoding collective, Wellcome Sanger Institute Tree of Life programme, et al. The genome sequence of *Aplidium turbinatum* (Savigny 1816), a colonial sea squirt. Wellcome Open Res. 2022;7:106.

45. Hirose E, Lopez JV, Oatley G, Sinclair E, Aunin E, Gettle N, et al. The chromosomal genome sequence of the photosymbiotic ascidian, Trididemnum clinides Kott, 1977 and its associated microbial metagenome sequences. Wellcome Open Res. 2025;10:357.

46. Bishop J, Wood C, Mrowicki RJ, Harley J, Marine Biological Association Genome Acquisition Lab, Darwin Tree of Life Barcoding collective, et al. The genome sequence of the Orange-tipped sea squirt, Corella eumyota Traustedt, 1882. Wellcome Open Res. 2024;9:146.

47. DeBiasse MB, Colgan WN, Harris L, Davidson B, Ryan JF. Inferring Tunicate Relationships and the Evolution of the Tunicate Hox Cluster with the Genome of Corella inflata. Genome Biol Evol. 2020;12:948–64.

48. Bishop J, Wood C, Marine Biological Association Genome Acquisition Lab, Darwin Tree of Life Barcoding collective, Wellcome Sanger Institute Tree of Life Management, Samples and Laboratory team, Wellcome Sanger Institute Scientific Operations: Sequencing Operations, et al. The genome sequence of a solitary sea squirt, Ascidia mentula (Müller, 1776). Wellcome Open Res. 2023;8:583.

49. Galià-Camps C, Schell T, Pegueroles C, Baranski D, Hamadou AB, Turon X, et al. Genomic richness enables worldwide invasive success. Research Square. 2024. 10.21203/rs.3.rs-3902873/v1.

50. Wei J, Zhang J, Lu Q, Ren P, Guo X, Wang J, et al. Genomic basis of environmental adaptation in the leathery sea squirt (Styela clava). Mol Ecol Resour. 2020;20:1414–31.

51. De Thier O, Tawfeeq MM, Faure R, Lebel M, Dru P, Blanchoud S, et al. First chromosome-level genome assembly of the colonial tunicate Botryllus schlosseri. bioRxiv. 2024. 10.1101/2024.05.29.594498.

52. Parra-Rincón E, Velandia-Huerto CA, Gittenberger A, Fallmann J, Gatter T, Brown FD, et al. The Genome of the “Sea Vomit” Didemnum vexillum. Life. 2021;11. 10.3390/life11121377.

53. Gabriel L, Brůna T, Hoff KJ, Ebel M, Lomsadze A, Borodovsky M, et al. BRAKER3: Fully automated genome annotation using RNA-seq and protein evidence with GeneMark-ETP, AUGUSTUS, and TSEBRA. Genome Res. 2024;34:769–77.

54. Emms DM, Kelly S. OrthoFinder: phylogenetic orthology inference for comparative genomics. Genome Biol. 2019;20:238.

55. Maeda T, Takahashi S, Yoshida T, Shimamura S, Takaki Y, Nagai Y, et al. Chloroplast acquisition without the gene transfer in kleptoplastic sea slugs, Plakobranchus ocellatus. Elife. 2021;10. 10.7554/eLife.60176.

56. Price MN, Dehal PS, Arkin AP. FastTree 2--approximately maximum-likelihood trees for large alignments. PLoS One. 2010;5:e9490.

57. Emms DM, Kelly S. STAG: Species tree inference from all genes. bioRxiv. 2018;:267914. 10.1101/267914.

58. Capella-Gutiérrez S, Silla-Martínez JM, Gabaldón T. trimAl: a tool for automated alignment trimming in large-scale phylogenetic analyses. Bioinformatics. 2009;25:1972–3.

59. Katoh K, Standley DM. MAFFT multiple sequence alignment software version 7: improvements in performance and usability. Mol Biol Evol. 2013;30:772–80.

60. Minh BQ, Schmidt HA, Chernomor O, Schrempf D, Woodhams MD, von Haeseler A, et al. IQ-TREE 2: New models and efficient methods for phylogenetic inference in the genomic era. Mol Biol Evol. 2020;37:1530–4.

61. Sanderson MJ. R8s: Inferring absolute rates of molecular evolution and divergence times in the absence of a molecular clock. Bioinformatics. 2003;19:301–2.

62. Han MV, Thomas GWC, Lugo-Martinez J, Hahn MW. Estimating gene gain and loss rates in the presence of error in genome assembly and annotation using CAFE 3. Mol Biol Evol. 2013;30:1987–97.

63. Wu T, Hu E, Xu S, Chen M, Guo P, Dai Z, et al. clusterProfiler 4.0: A universal enrichment tool for interpreting omics data. Innovation (Camb). 2021;2:100141.

64. Wickham H. Ggplot2: Elegant graphics for data analysis. 1st edition. New York, NY: Springer; 2009.

65. Rice P, Longden I, Bleasby A. EMBOSS: The European molecular biology open software suite. Trends Genet. 2000;16:276–7.

66. Wright F. The “effective number of codons” used in a gene. Gene. 1990;87:23–9.

67. Wang Y, Jiang D, Guo K, Zhao L, Meng F, Xiao J, et al. Comparative analysis of codon usage patterns in chloroplast genomes of ten Epimedium species. BMC Genom Data. 2023;24:3.

68. Li B, Dewey CN. RSEM: accurate transcript quantification from RNA-Seq data with or without a reference genome. BMC Bioinformatics. 2011;12:323.

69. Diesh C, Stevens GJ, Xie P, De Jesus Martinez T, Hershberg EA, Leung A, et al. JBrowse 2: a modular genome browser with views of synteny and structural variation. Genome Biol. 2023;24:74.

70. Plessy C, Mansfield MJ, Bliznina A, Masunaga A, West C, Tan Y, et al. Extreme genome scrambling in marine planktonic Oikopleura dioica cryptic species. Genome Res. 2024;34:426–40.

71. Kocot KM, Tassia MG, Halanych KM, Swalla BJ. Phylogenomics offers resolution of major tunicate relationships. Mol Phylogenet Evol. 2018;121:166–73.

72. Shenkar N, Koplovitz G, Dray L, Gissi C, Huchon D. Back to solitude: Solving the phylogenetic position of the Diazonidae using molecular and developmental characters. Mol Phylogenet Evol. 2016;100:51–6.

73. Elsik CG, Worley KC, Bennett AK, Beye M, Camara F, Childers CP, et al. Finding the missing honey bee genes: lessons learned from a genome upgrade. BMC Genomics. 2014;15:86.

74. Elliott TA, Gregory TR. What’s in a genome? The C-value enigma and the evolution of eukaryotic genome content. Philos Trans R Soc Lond B Biol Sci. 2015;370:20140331.

75. Anselmi C, Ishizuka KJ, Palmeri KJ, Burighel P, Voskoboynik A, Hotta K, et al. Speed vs completeness: a comparative study of solitary and colonial tunicate embryogenesis. Front Cell Dev Biol. 2025;13:1540212.

76. Delsuc F, Brinkmann H, Chourrout D, Philippe H. Tunicates and not cephalochordates are the closest living relatives of vertebrates. Nature. 2006;439:965–8.

77. Tsagkogeorga G, Turon X, Hopcroft RR, Tilak M-K, Feldstein T, Shenkar N, et al. An updated 18S rRNA phylogeny of tunicates based on mixture and secondary structure models. BMC Evol Biol. 2009;9:187.

78. Shenkar N, Swalla BJ. Global diversity of Ascidiacea. PLoS One. 2011;6:e20657.

79. Smith ML, Hahn MW. New approaches for inferring phylogenies in the presence of paralogs. Trends Genet. 2021;37:174–87.

80. Inoue J, Satoh N. ORTHOSCOPE: An automatic web tool for phylogenetically inferring bilaterian orthogroups with user-selected taxa. Mol Biol Evol. 2019;36:621–31.

81. Kyriacou RG, Mulhair PO, Holland PWH. GC content across insect genomes: Phylogenetic patterns, causes and consequences. J Mol Evol. 2024;92:138–52.

82. Šmarda P, Bureš P, Horová L, Leitch IJ, Mucina L, Pacini E, et al. Ecological and evolutionary significance of genomic GC content diversity in monocots. Proc Natl Acad Sci U S A. 2014;111:E4096–102.

83. Romiguier J, Ranwez V, Douzery EJP, Galtier N. Contrasting GC-content dynamics across 33 mammalian genomes: relationship with life-history traits and chromosome sizes. Genome Res. 2010;20:1001–9.

84. Bernardi G, Bernardi G. Compositional constraints and genome evolution. J Mol Evol. 1986;24:1–11.

85. Clessin A, Joseph J, Lartillot N. Evolution of GC-biased gene conversion by natural selection. Genetics. 2025;230:iyaf111.

86. Romiguier J, Roux C. Analytical biases associated with GC-content in molecular evolution. Front Genet. 2017;8:16.

87. Glémin S, Arndt PF, Messer PW, Petrov D, Galtier N, Duret L. Quantification of GC-biased gene conversion in the human genome. Genome Res. 2015;25:1215–28.

88. Deng W, Henriet S, Chourrout D. Prevalence of mutation-prone microhomology-mediated end joining in a chordate lacking the c-NHEJ DNA repair pathway. Curr Biol. 2018;28:3337–3341.e4.

89. Moggioli G, Panossian B, Sun Y, Thiel D, Martín-Zamora FM, Tran M, et al. Distinct genomic routes underlie transitions to specialised symbiotic lifestyles in deep-sea annelid worms. Nat Commun. 2023;14:2814.

90. Krokan HE, Bjørås M. Base excision repair. Cold Spring Harb Perspect Biol. 2013;5:a012583.

91. Shito TT, Oka K, Hotta K. Multimodal factor evaluation system for organismal transparency by hyperspectral imaging. PLoS One. 2023;18:e0292524.

92. Hotta K, Miyasaka SO, Oka K, Shito TT. Staring into a crystal ball: understanding evolution and development of in vivo aquatic organismal transparency. Front Ecol Evol. 2024;12:1428976.

93. Sinha RP, Häder DP. UV-induced DNA damage and repair: a review. Photochem Photobiol Sci. 2002;1:225–36.

