## Supplemental Data 2 for "Genomic resources of *Ascidiella aspersa* and comparative analysis across tunicates reveal class-level features and evolutionary diversification"

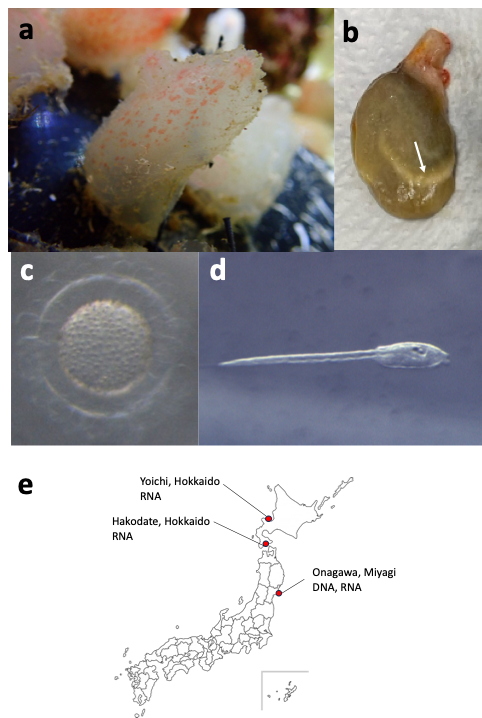

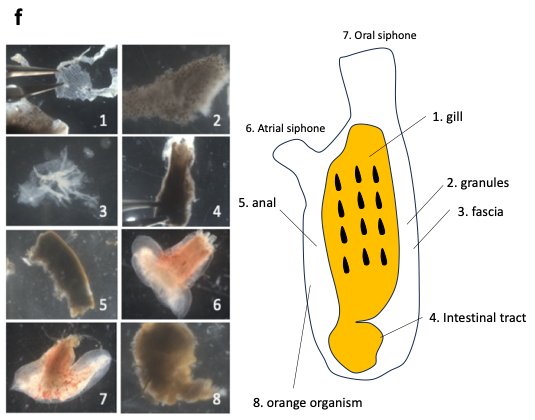

**Fig. S1 The adult and embryos of Ascidiella aspersa**

a) Pictures of adult *A. aspersa*. b) Adults excluding tunics. Arrowhead show the sperm duct. c) *A. aspersa* eggs and d) larvae with high transparency. e) Adult organs from which RNA was extracted. f) Sampling sites of *A. aspersa*.

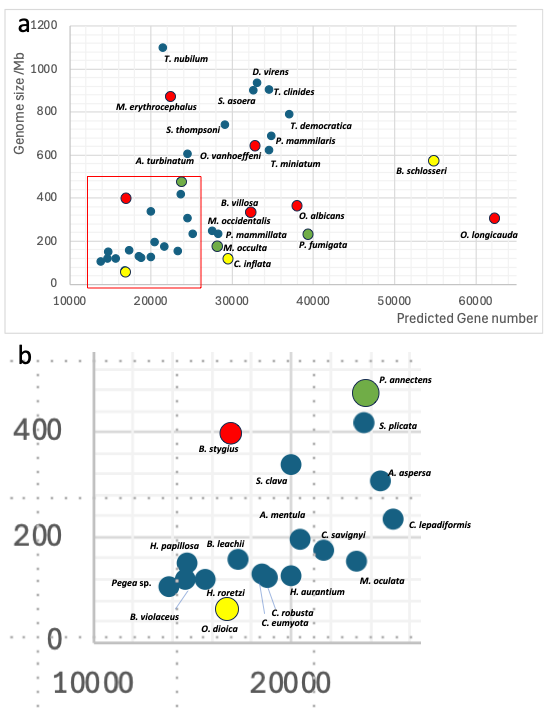

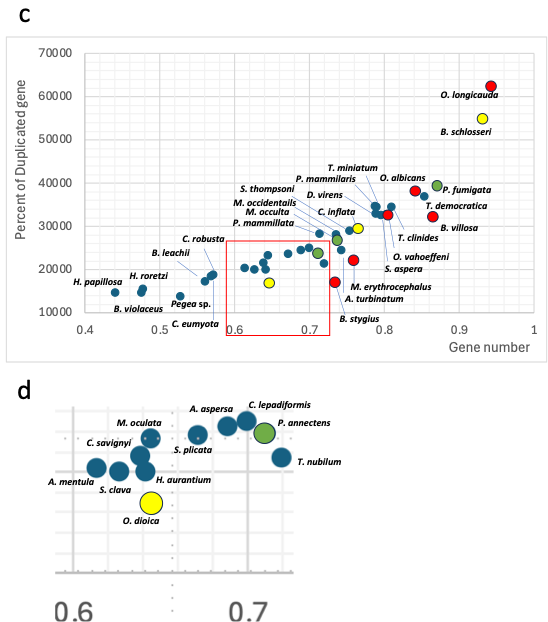

**Fig. S2 Gene numbers and genome sizes**

Scatter plot of genome size and predicted gene numbers of 39 tunicates. Red dots indicate tunicate genomes with BUSCO scores <80%. Yellow dots indicate scores of 85%, and green dots indicate scores <90%, respectively. a) All species. b) Close-up view of the orange rectangle in a). c) Duplicated gene ratio to whole gene numbers. d) Close-up view of c).

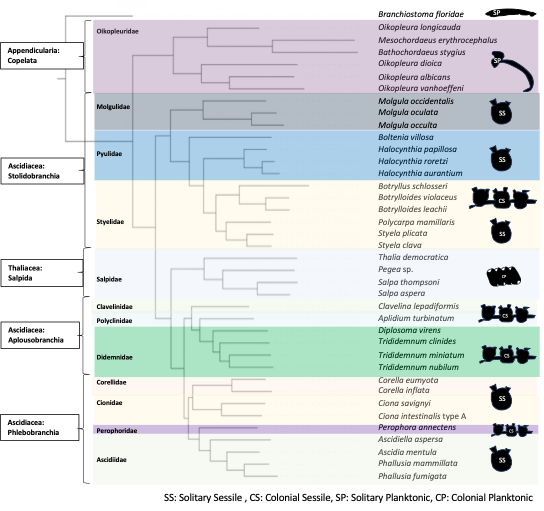

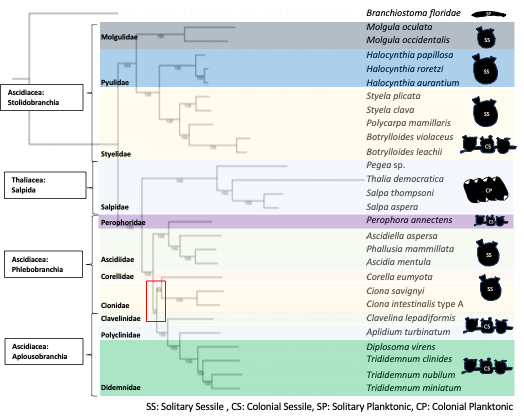

**Fig. S3 Phylogenetic tree**

a) The species tree with 39 species generated by STAG in OrthoFinder. The support values at internal bipartitions represent the proportion of input gene trees that support each bipartition. b) Phylogenetic tree constructed with ML analysis using 28 tunicates genome with BUSCO scores higher than approximately 90%. Nodes in the red square show 94% ultra bootstrap.

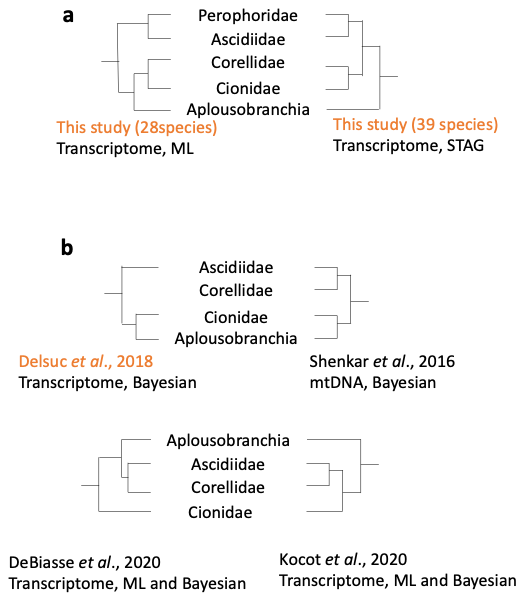

**Fig. S4 Phylogenetic tree compared to recent works**

Comparison of phylogenetic trees in this study and those in four recent literatures. Dataset types and methods of phylogenetic analysis are described. Delsuc et al. 2018 is the only study consistent with our tree using 28 species.

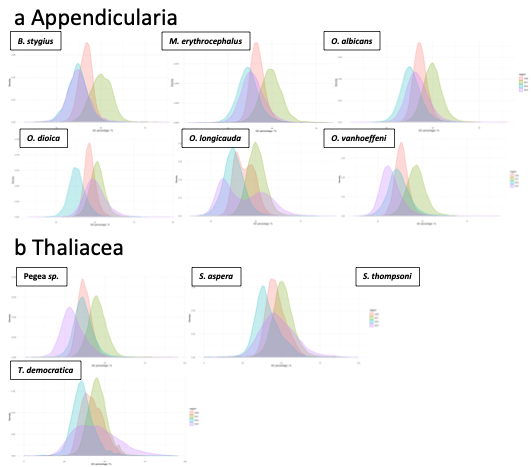

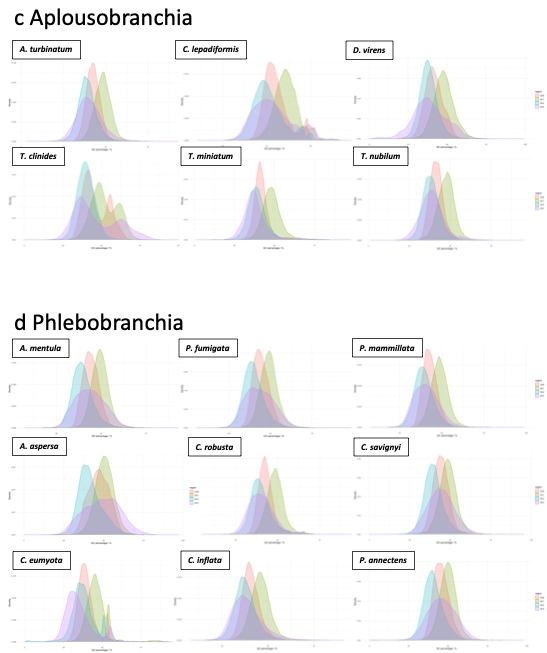

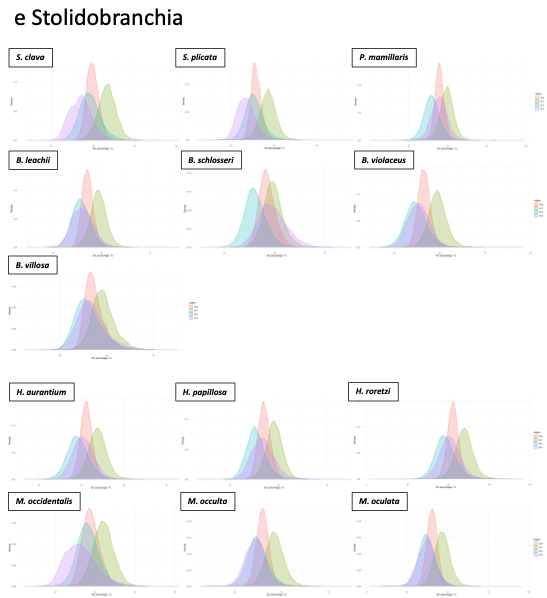

**Fig. S5 GC content of 38 tunicates in different codon regions**

Density plot of GC content in all genes and GC contents. Red, green, blue, and purple colors represent whole CDS, first codon, second codon, and third codon, respectively.

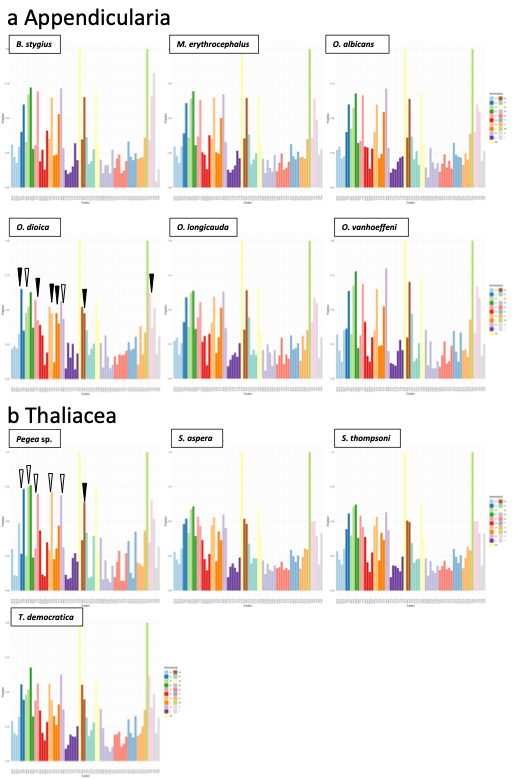

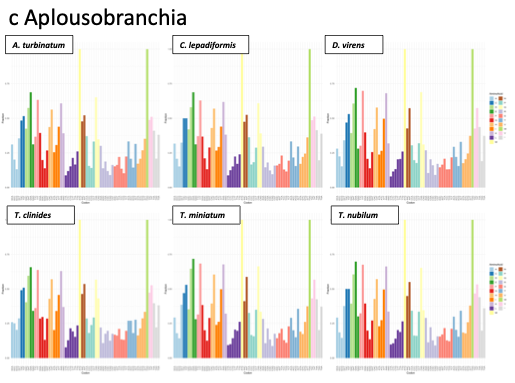

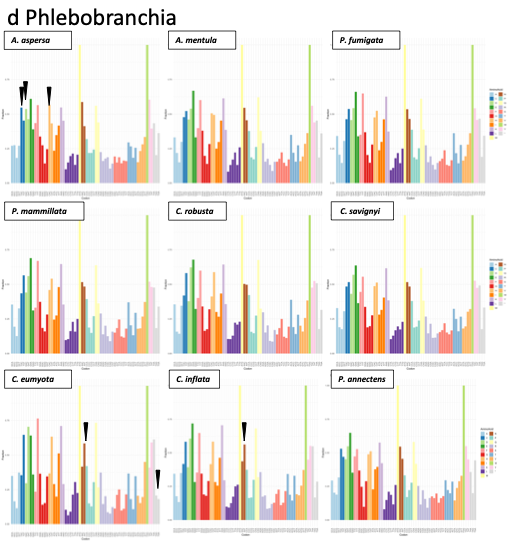

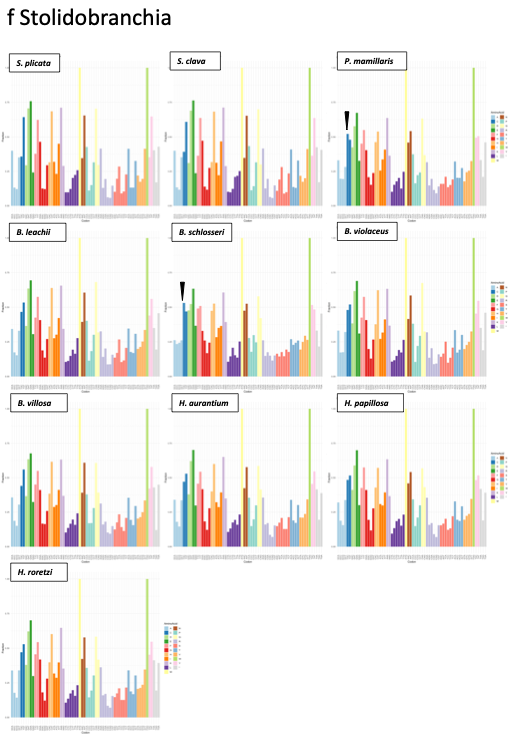

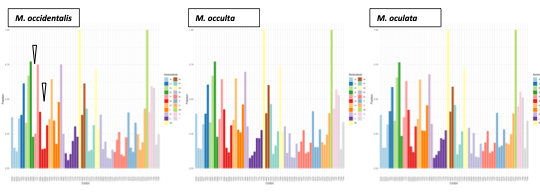

**Fig. S6 Codon usage in 38 tunicates**

Codon usages in 38 tunicates. Different colors represent different amino acids. Black pins indicate the reversal of codon usage compared to other Ascididae species. White pins indicate codon usage changes that are not as significant as reversals.
