## Supplemental Data 3 for "Genomic resources of *Ascidiella aspersa* and comparative analysis across tunicates reveal class-level features and evolutionary diversification"

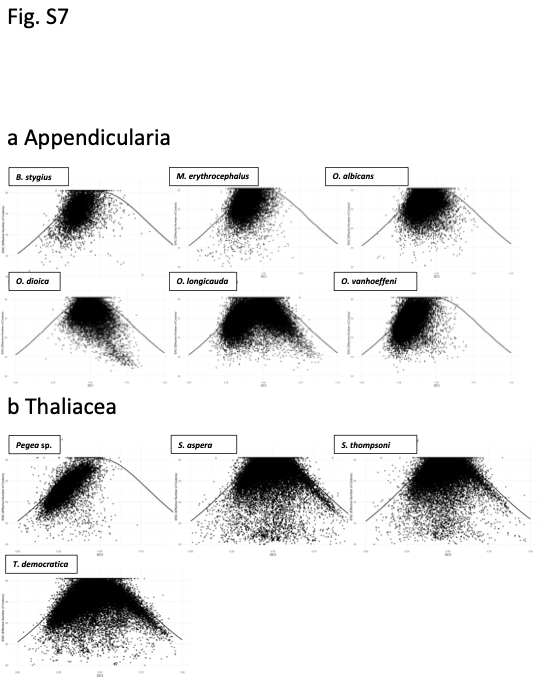

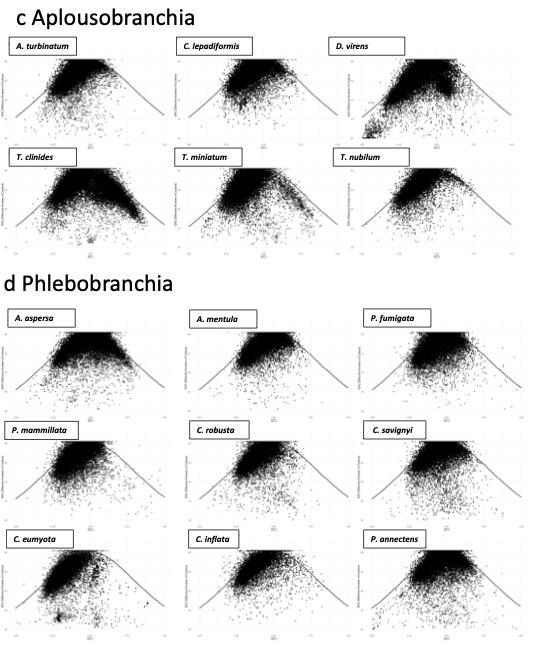

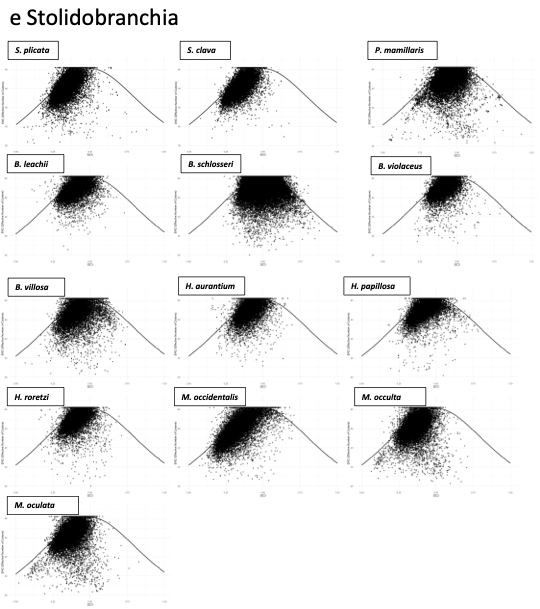

**Fig. S7 ENC-GC3 plot of 38 tunicates**

ENC-GC3 plot of 38 tunicates.

a: Annotation Page

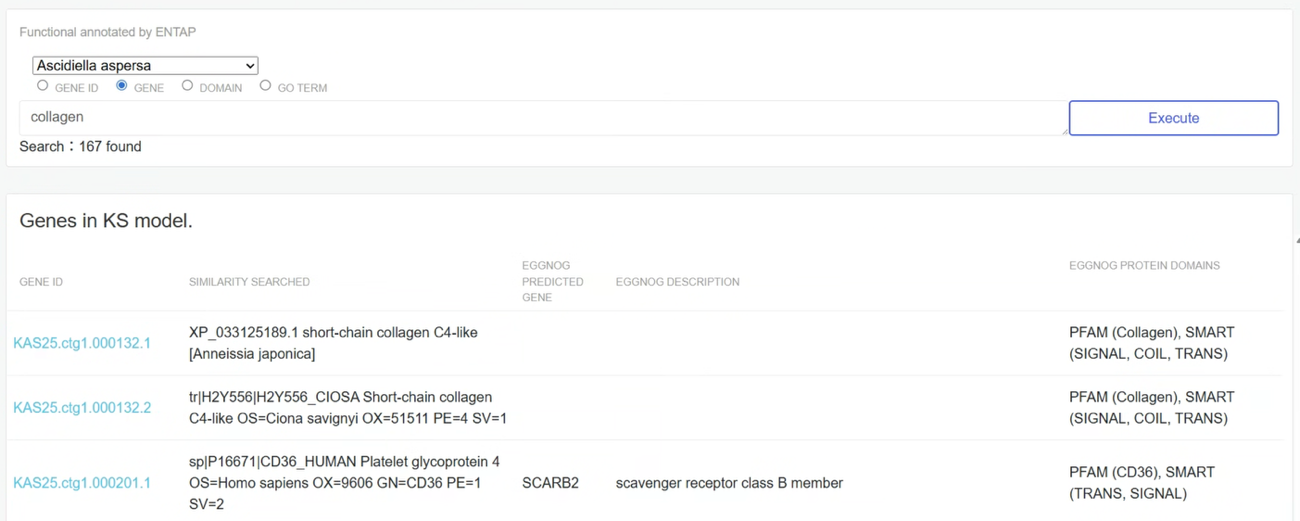

b: Individual gene

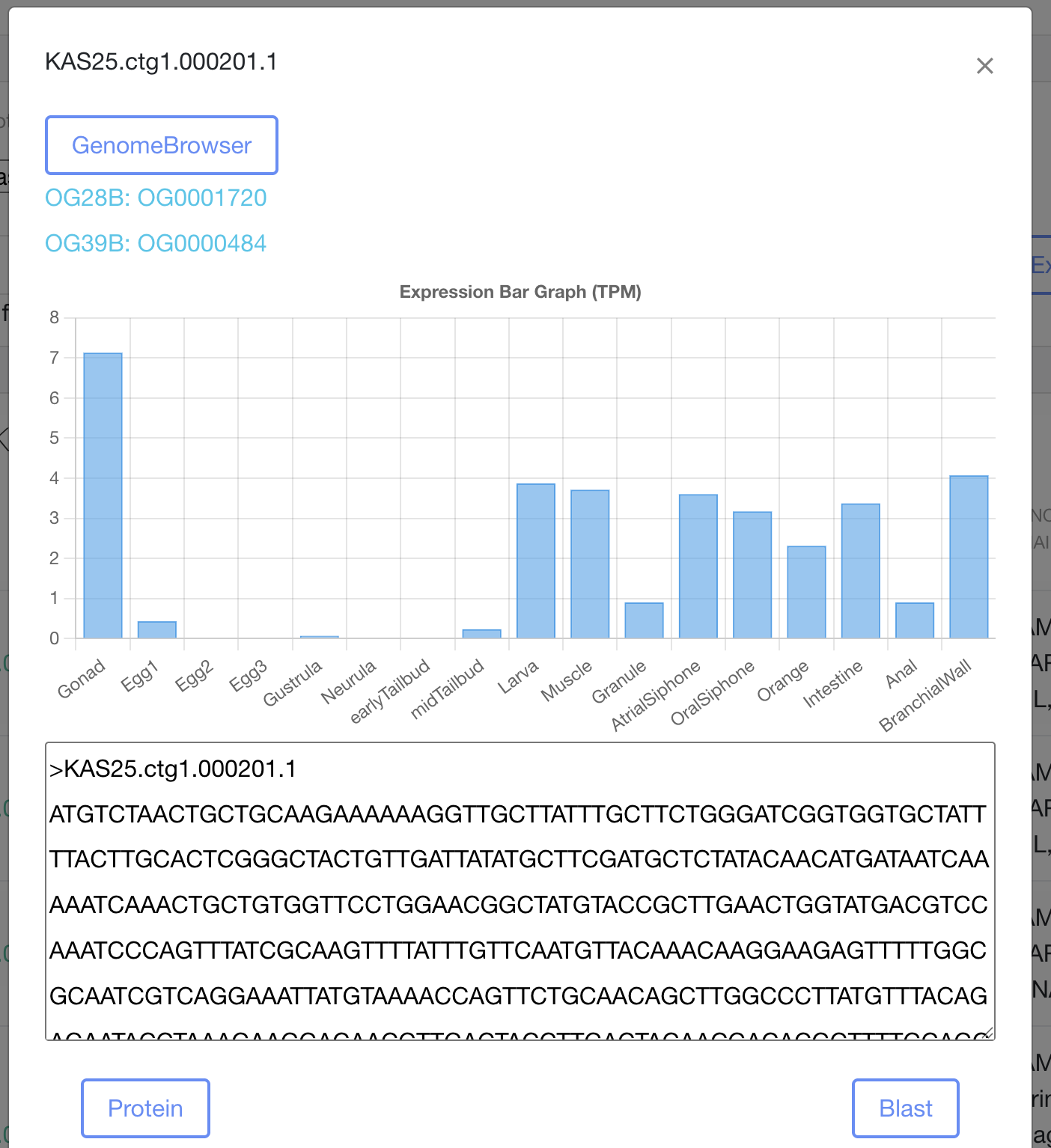

c: genome browser

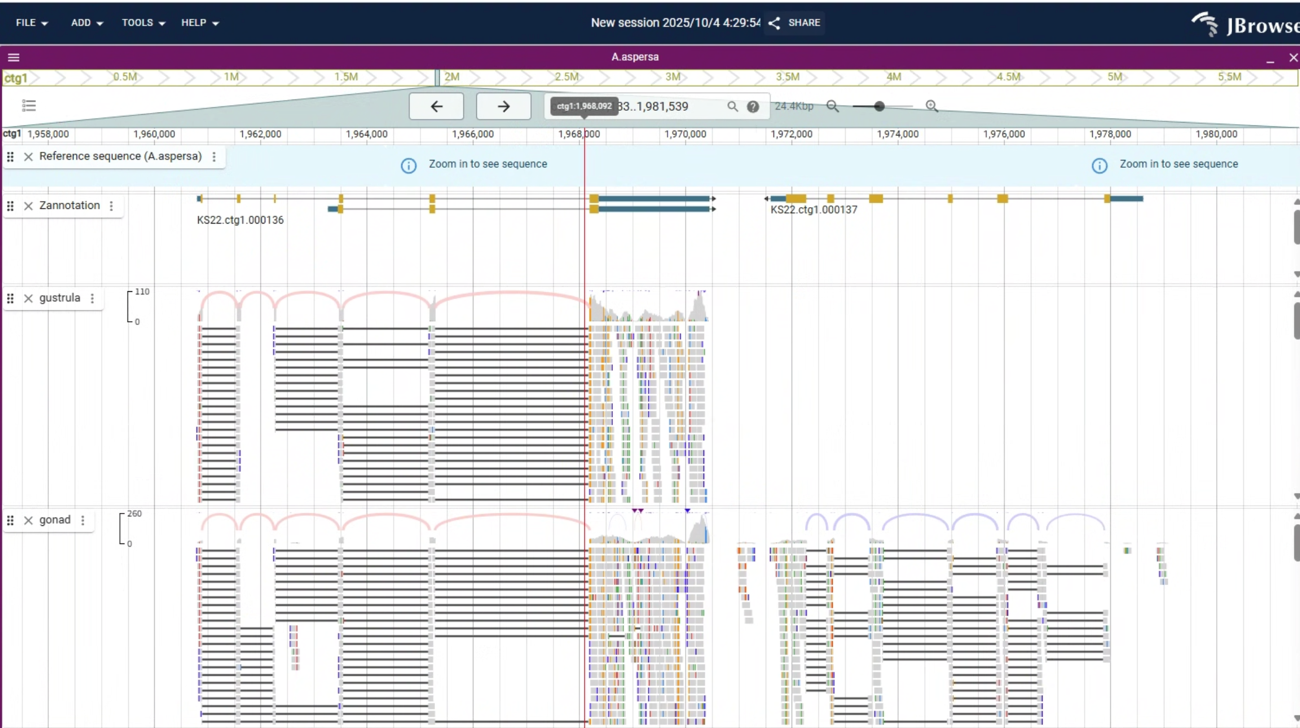

d: Phylogram of individual gene

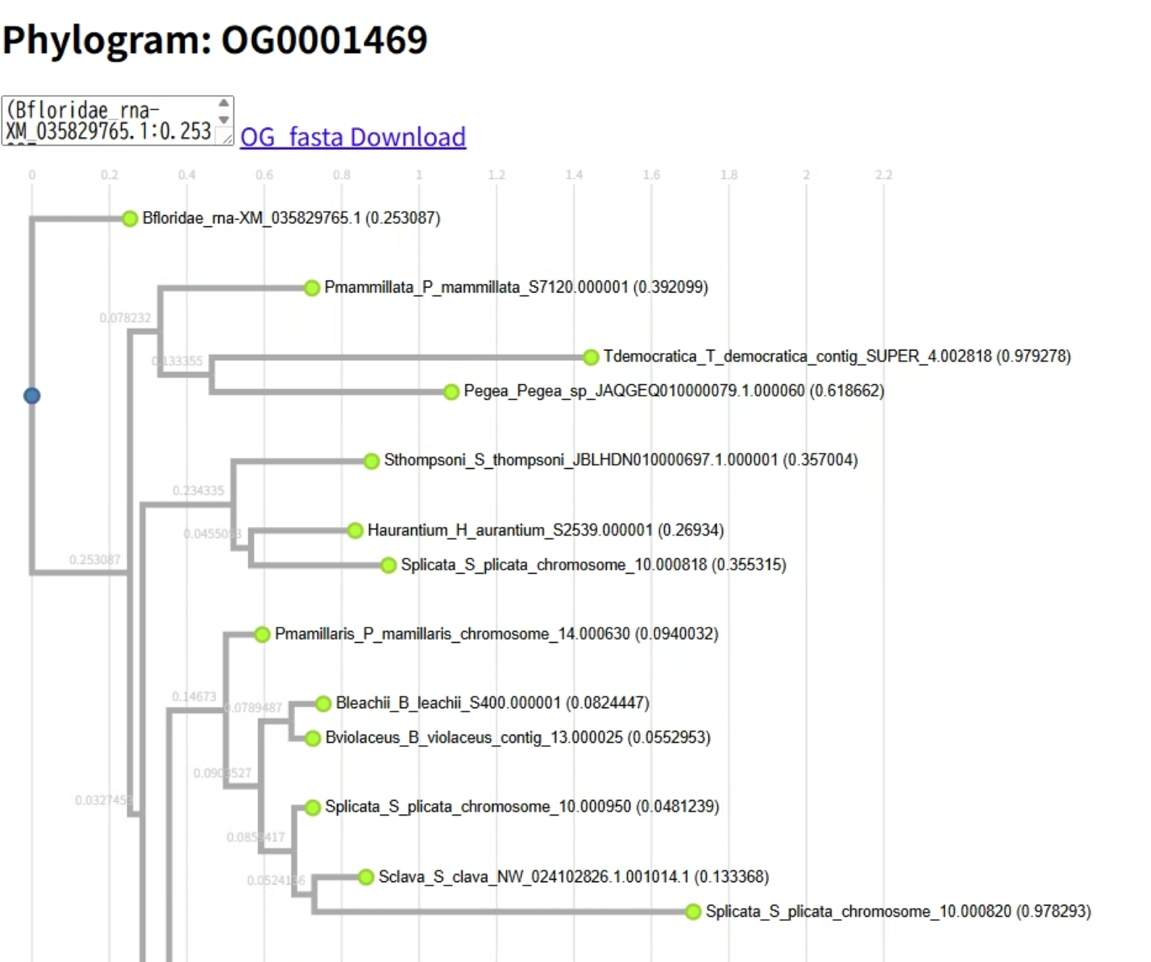

e: Blast page

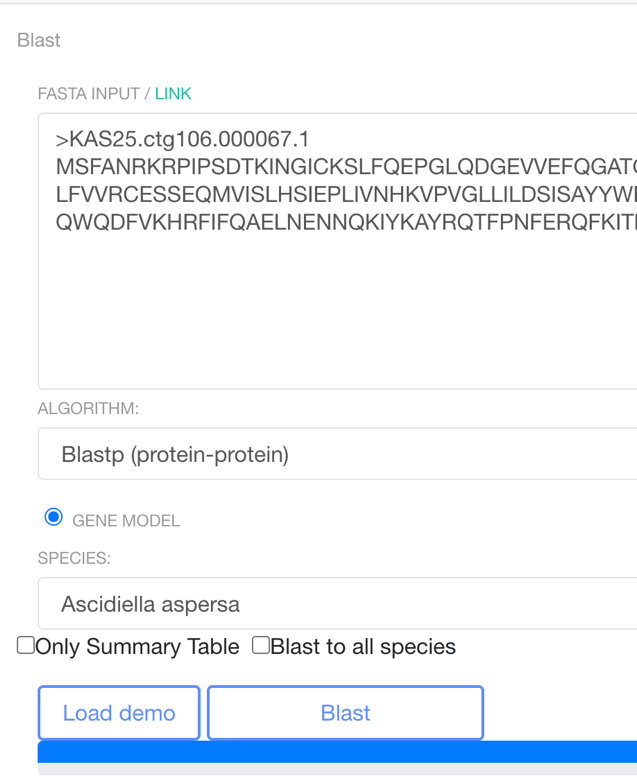

f: Blast result page

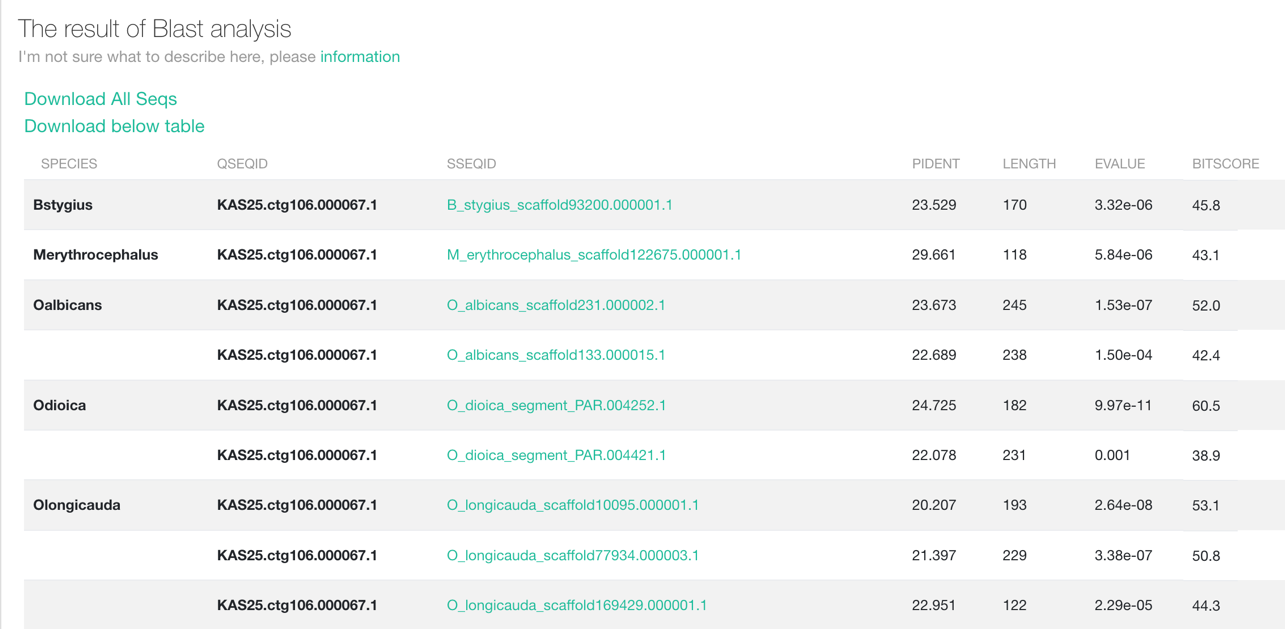

**Fig. S8 Screens of the TUNOME database**

a) Annotation browser constructed using functional annotations derived from ENTAP. b) Individual gene view. Annotations, bar graphs of transcripts per million (TPM), and sequences can be viewed. d) Genome browser of 38 species. Direct access to individual genes from b). d) Specific gene tree of an orthogroup. Orthogroups of 28 species and 39 species were uploaded. The Newick tree and FASTA file can also be downloaded. e) BLAST page allows searches against genomes and gene models from 38 tunicates. A search to all species can be performed by checking a box. f) BLAST result of the search across all species. All hit sequences and table can be downloaded.

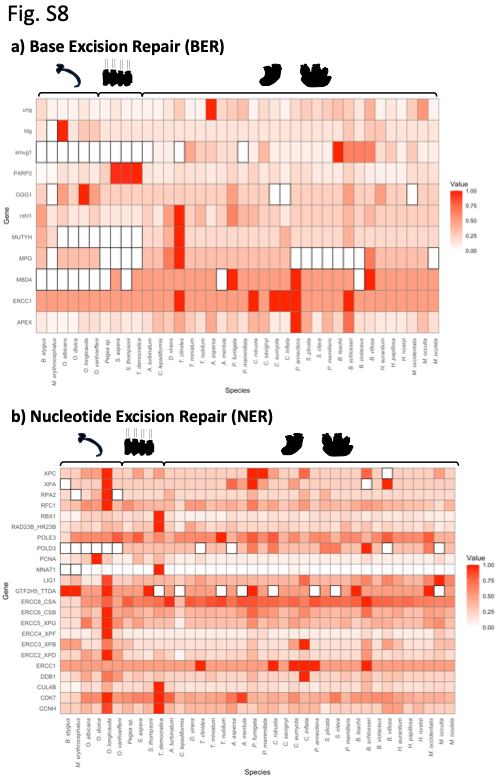

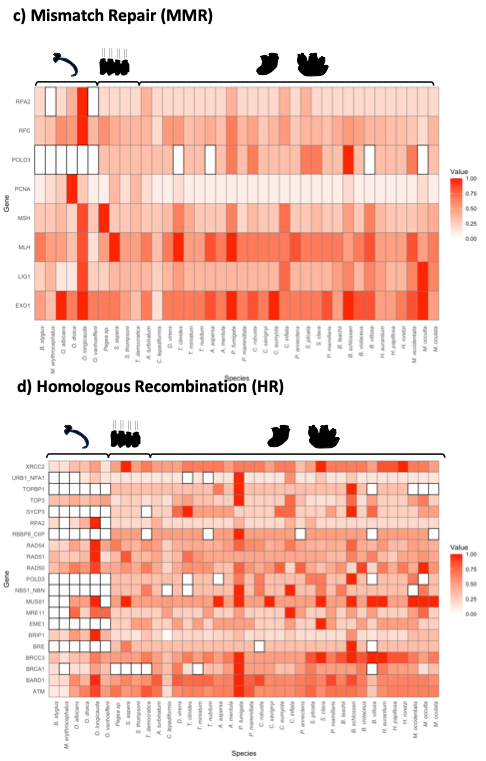

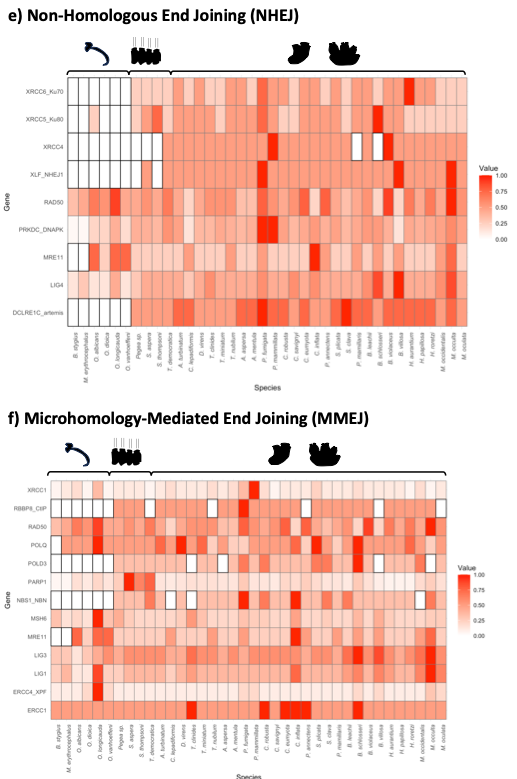

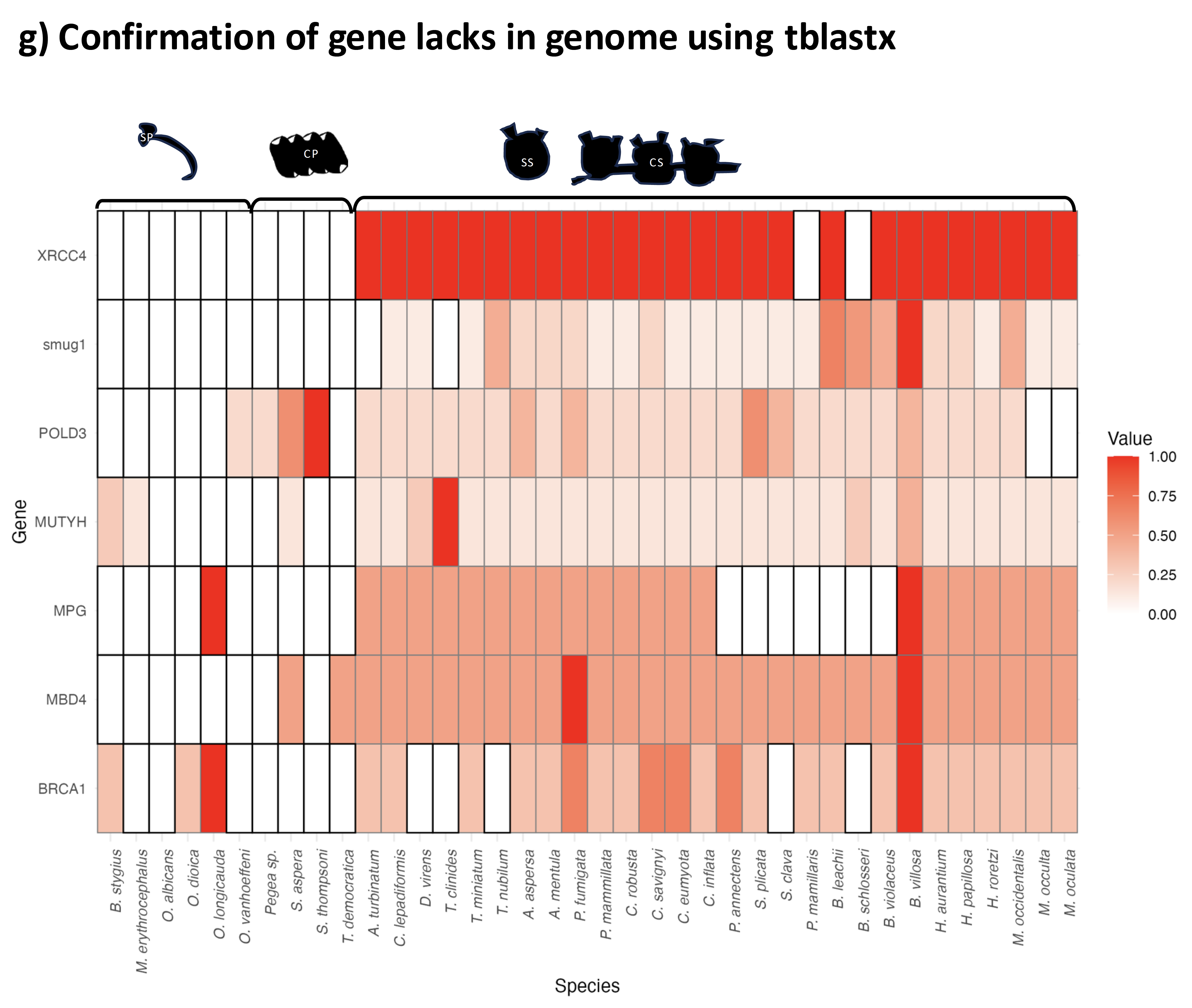

**Fig. S9 DNA repair genes**

DNA repair genes in tunicates. Individual genes were blasted against all tunicate protein databases via the Tunome database with a cutoff e-value <0.005. Hit numbers were converted to relative values based on the highest hit number among species. Absent genes were highlighted with surrounding black lines. Genes associated with a) base excision repair, b) nucleotide excision repair, c) mismatch repair, d) homologous recombination, e) non-homologous end joining, and f) microhomology-mediated end joining were selected according to Moggioli et al. 2023. Appendicularia, Thaliacea, and Styelidae species show shared absences of specific repair genes. g) verifying by searching tblastx to genomes. Gene lacks observed in above were verified at genome level.

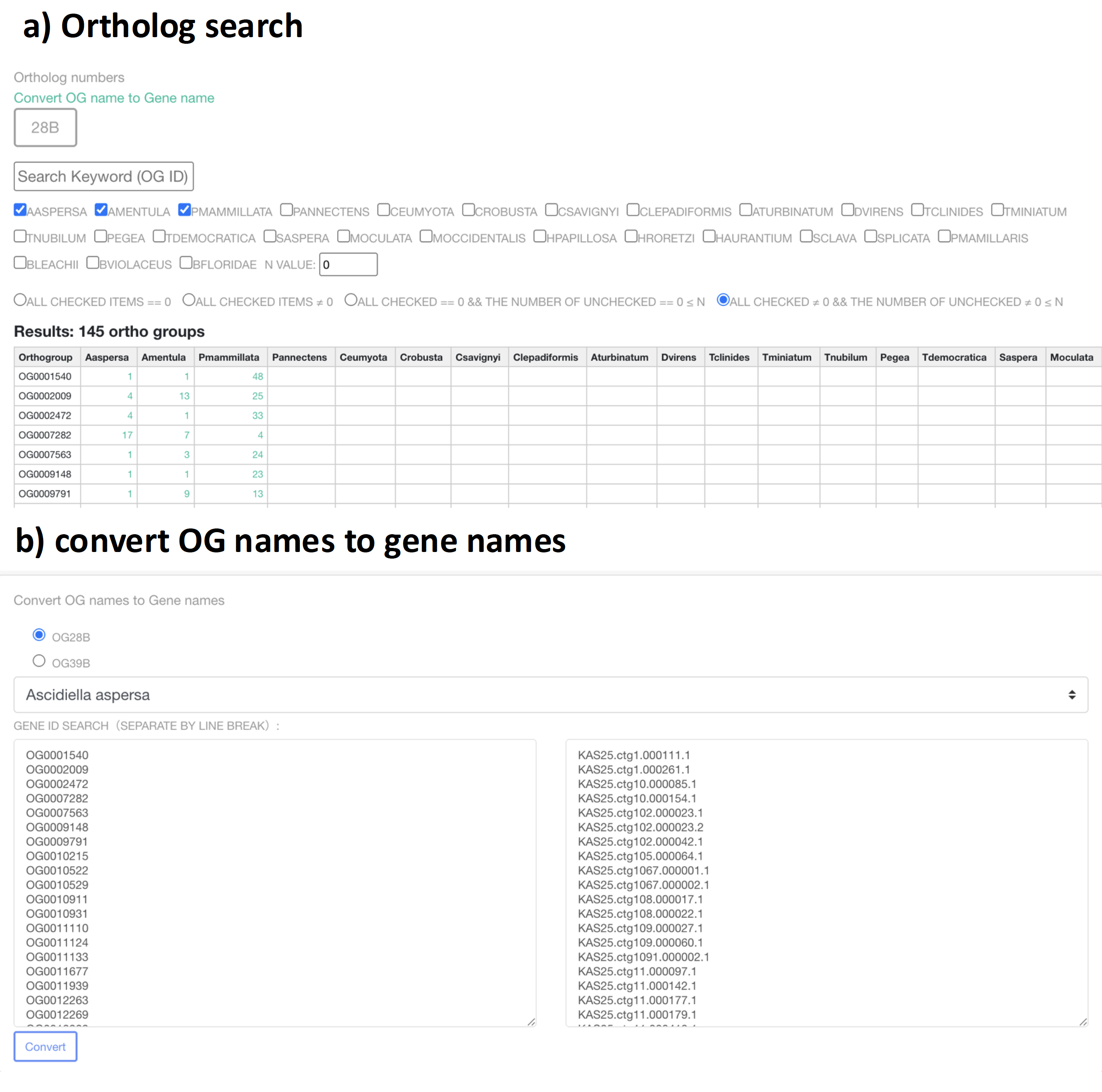

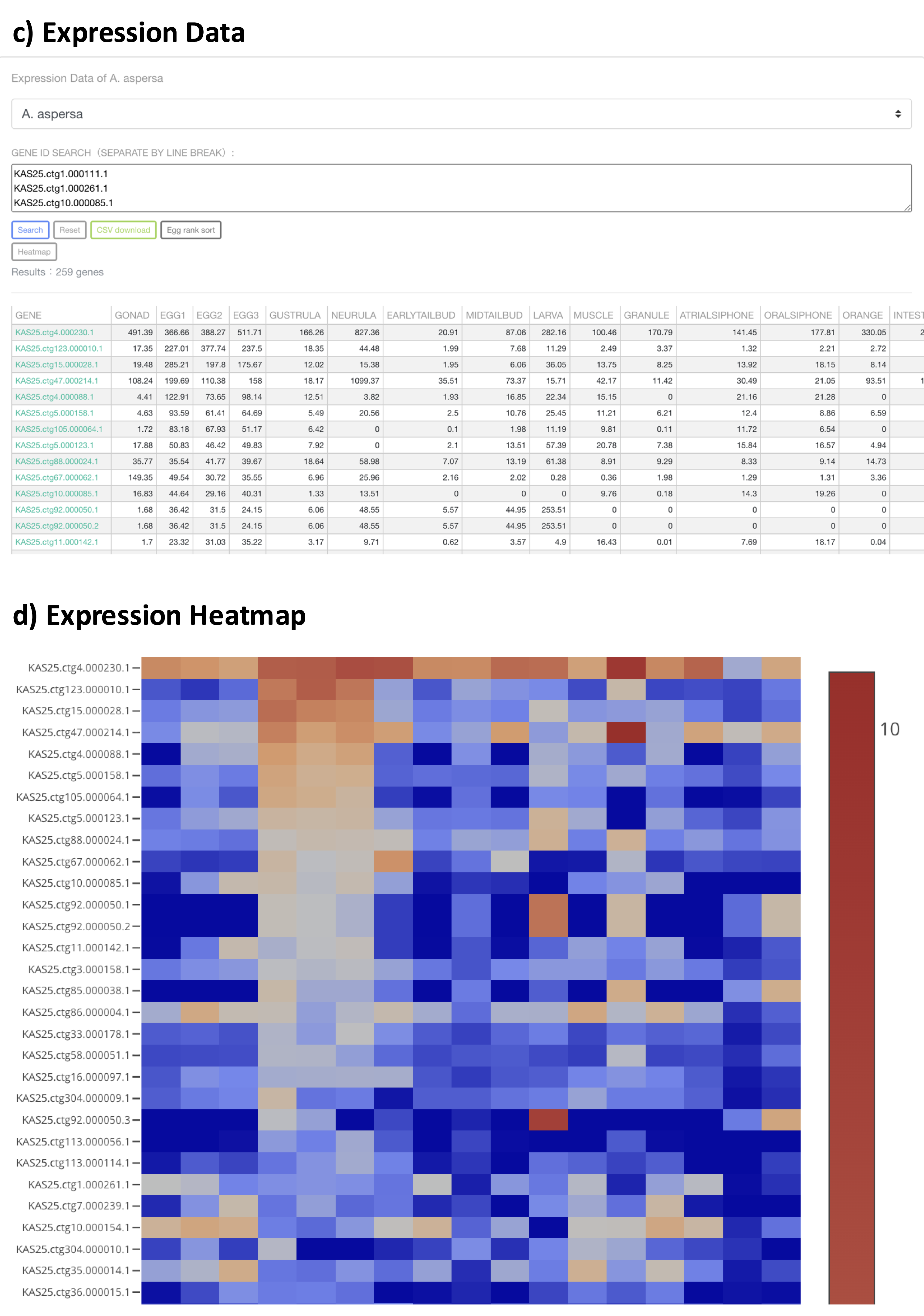

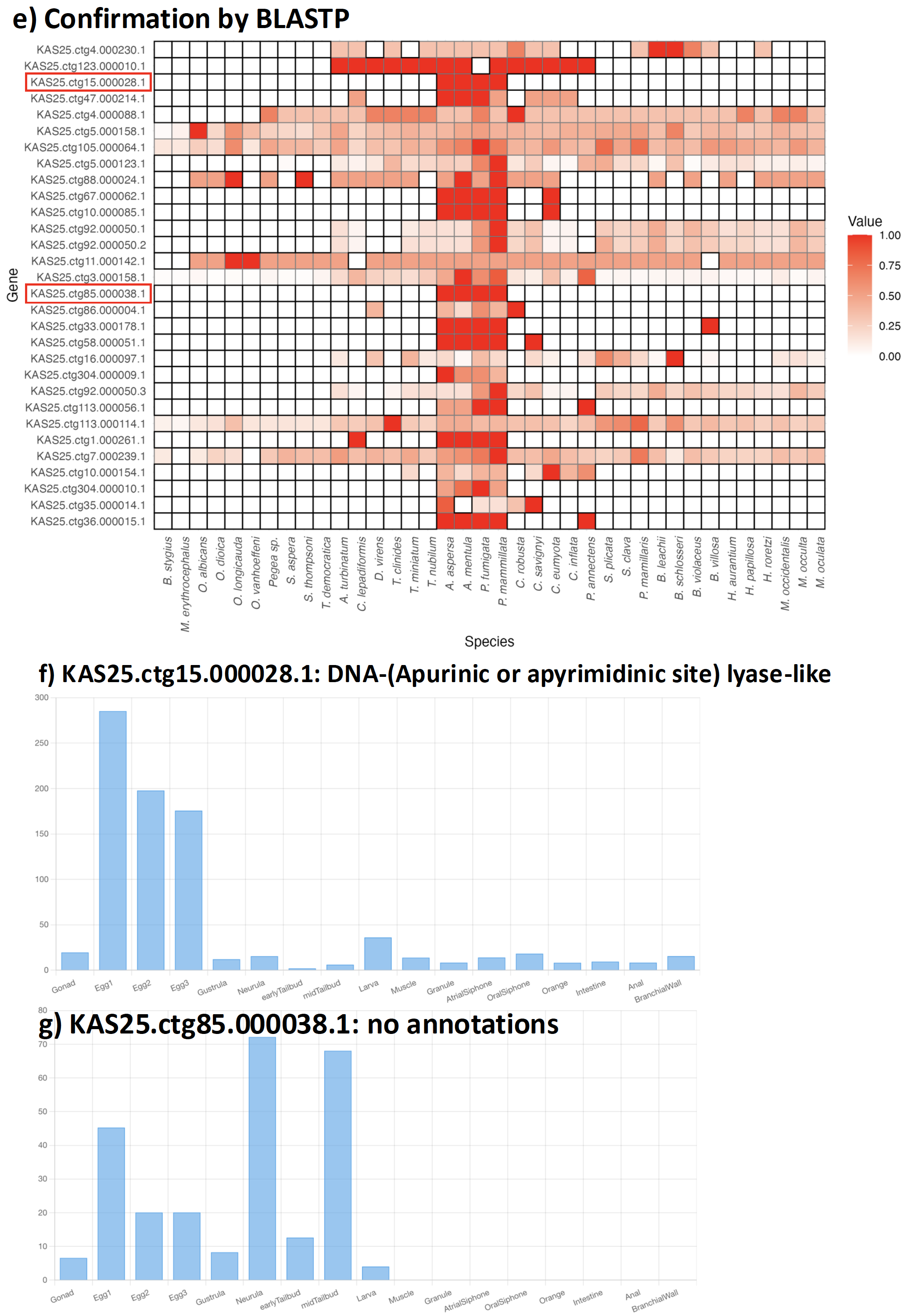

**Fig. S10 Exploring ascidiids-specific orthogroups using TUNOME**

a) Ortholog search view. After selecting species of interest at the top of the page, candidate orthogroups identified using OrthoFinder v2 are filtered according to the conditions specified below. The filtering options are defined as follows: (i) “all checked = 0” indicates that none of the selected species contain the orthogroup; (ii) “all checked ≠ 0” indicates that all selected species share the orthogroup; (iii) “all checked = 0 and the number of unchecked species ≤ N” indicates that none of the selected species contain the orthogroup, while at most N unselected species do; and (iv) “all checked ≠ 0 and the number of unchecked species ≤ N” indicates that all selected species share the orthogroup, while at most N unselected species also share it. For identification of ascidiids-specific genes, select *A. aspersa*, *A. mentula*, and *P. mammillata*, choose condition (iv), and set N = 0. b) Convert OG names to gene names page linked from a). You can paste specific orthogroups names copied from the page of a), and select species name, then corresponding genes in selected species, contained in the orthogroups were obtained. c) Expression data view. You can paste gene names of *A. aspersa* obtained from the page of b), expression data (TPM obtained by RSEM) were viewed. Press the egg sort button and Heatmap button. the expression data were visualized in the bottom of the page d). For the confirmation of this orthogroup analysis, we confirmed by performing Blastp in TUNOME and visualized Rscripts. In top 30 genes, two genes were confirmed as ascidiids-specific e), and expressions were individually displayed f)g).

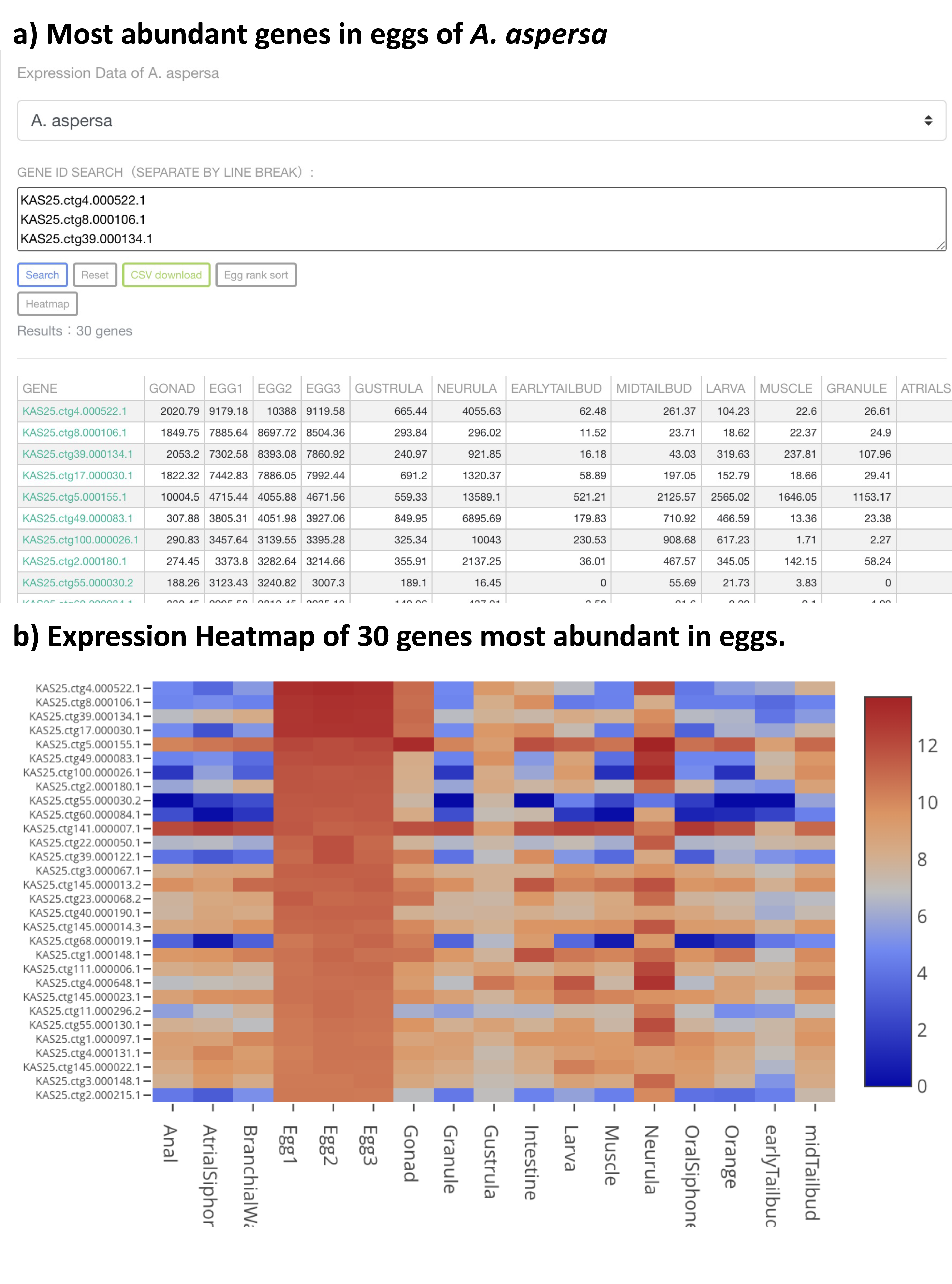

**Fig. S11 Exploring most abundant genes in eggs transcriptome of *A. aspersa***

a) To identify the most abundant genes in eggs of A. aspersa, the “eggs sort” button was selected in the expression data view, and the top 30 genes were retrieved. These gene names were entered into the exploration box, followed by clicking the “Explore” button and then the “Heatmap” button, resulting in a heatmap displayed at the bottom of the page b).
